# Loss of neurofibromin accelerates uveal and dermal melanoma formation driven by GNAQ

**DOI:** 10.1101/2024.06.26.600903

**Authors:** Anne Nathalie Longakit, Oscar Urtatiz, Amy Luty, Christina Zhang, Chloe Hess, Alyssa Yoo, Hannah Bourget, Catherine D. Van Raamsdonk

## Abstract

Neurofibromin is a very large and complex tumor suppressor, whose loss can synergize with other MAPK pathway mutations to promote melanoma in the skin. In this paper, we investigated whether *NF1* loss has a role in other melanomas, such as those that form in the dermis or eye (uveal tract). We found that heterozygous 17q11.2 loss that includes the *NF1* locus is an uncommon, but recurrent phenomenon in human dermal and uveal melanomas described previously. We studied the effects of *Nf1* haploinsufficiency in mice expressing oncogenic GNAQ^Q209L^ in melanocytes and Schwann cells of peripheral nerves using the *Plp1-creERT* transgene, with tamoxifen given at 5 weeks of age. *Nf1* haploinsufficiency accelerated dermal and uveal melanoma formation. We studied the effects of *Nf1* loss in these melanomas using RNAseq. Many of the differentially expressed genes were homologous to genes whose expression correlates with prognosis in human uveal melanoma. Of particular interest was the up-regulation of cAMP signaling and its connection to protein kinase A, which is mutant in malignant melanotic nerve sheath tumors (MMNSTs). An unexpected finding was that oncogenic GNAQ was sufficient by itself to drive peripheral nerve sheath-like neoplasms in the mice. Hence, these studies reveal new insight into both melanocyte and Schwann cell transformation.

## INTRODUCTION

MAP Kinase (MAPK) pathway activation is one of the key events in melanomagenesis, as well as in many other cancers [1]. Somatic mutations that activate the MAPK pathway in common cutaneous melanoma include oncogenic mutations at specific hotspots in *NRAS* and *BRAF* [2,3] and tumor suppressor mutations in negative regulators, such as *RASA2* and *NF1* [4,5]. The heterotrimeric G protein alpha subunits, Gα_q_ and Gα_11_, also participate in MAPK activation, and oncogenic mutations in these two genes are very frequent in melanocytic lesions in the dermis, meninges of the central nervous system, and uveal tract of the eye [6,7,8,9]. The oncogenic mutations in *GNAQ* and *GNA11* at Q209 or R183 cause constitutive activity, preventing Gα_q_ and Gα_11_ from performing GTP hydrolysis and returning to an inactive state. Gα_q_ and Gα_11_ activate phospholipase C-beta 4, which stimulates protein kinase C by way of the second messenger, DAG. PKC activates RASGRP3 feeding into the MAPK pathway [10].

While oncogenic MAPK mutations are mutually exclusive with each other [11], the tumor suppressor *NF1* mutations have more of a cooperative or additive effect. For example, nearly two-thirds of *NF1* mutant cutaneous melanomas carry a second MAPK gene hit [5,12]. *NF1* mutations are present in 5% of *BRAF*, 13% of *NRAS* and 50% of *RASA2* mutant cutaneous melanomas. When *NF1* mutations occur in combination with another MAPK hit, the *NF1* mutations are more likely to be heterozygous, presumably acting through haploinsufficiency [4]. When *NF1* is the only mutated MAPK gene, both alleles of *NF1* are typically mutant. This occurs by either compound heterozygous mutations or one *NF1* focal mutation plus loss of heterozygosity. Zeng *et al* and others have suggested that MAPK pathway activation in melanoma is not an “all or nothing” phenomenon [13].

In this paper, we investigated a possible cooperative role of *NF1* loss in the context of *GNAQ* mutant melanoma. *NF1*, which is located at 17q11.2, encodes the very large neurofibromin protein [14]. Neurofibromin is a multifunctional protein that, in context specific ways, regulates MAPK, PI3K/AKT/mTOR, Rho/ROCK/LIMK2/cofilin, PKA-Ena/VASP and cAMP/PKA signaling (reviewed in [15].) This affects various cellular processes related to tumorigenesis, including proliferation, migration, cytoskeletal dynamics, and apoptosis.

In addition, heterozygous loss of function mutations in *NF1* cause neurofibromatosis type 1. In addition to many other symptoms, individuals with neurofibromatosis develop neurofibromas when there is a second, somatic hit in *NF1* in Schwann cells. Neurofibromas are complex tumors containing Schwann cells, fibroblasts, perineural cells, and mast cells in a variably myxoid background. People with neurofibromatosis type 1 also have pigmentation alterations, such as generalized and subtle skin hyper-pigmentation and flat, circumscribed Cafe au lait macules [16]. There are case reports of uveal melanoma occurring in patients with neurofibromatosis type 1 [17,18,19] and it has been estimated that twice as many cases of uveal melanoma have been reported than would be expected by chance [20]. Another study found that *NF1* expression was down regulated in uveal melanoma [21].

We previously studied conditional and constitutive knockout *Nf1* mutations in mice and found that there is tail skin hyper-pigmentation [22,23]. Histology of the mouse tail skin showed that the epidermis was darker, similar to the generalized skin hyper-pigmentation in human *NF1* patients. In addition, the dermis was hyper-pigmented in the *Nf1* mutant mice, suggesting that *NF1* regulates non-epidermal melanocytes as well [22].

To determine whether *NF1* loss has a role in GNAQ-driven melanoma, such as forms in the dermis or eye, we surveyed the published literature and the Cancer Genome Atlas (TCGA) uveal melanoma dataset (UVM) to see if there were copy number changes in *NF1* reported in intra-dermal melanocytic lesions (“blue nevus” types) or uveal melanoma. We found that 14% of malignant, but not benign, intra-dermal lesions exhibited copy number loss that included the *NF1* gene. There were also two cases of uveal melanoma with *NF1* copy number loss among the 80 TCGA UVM samples, and these cases had some other intriguing molecular features. Next, to test the effect of *Nf1* loss in a model system, we forced the expression of oncogenic GNAQ^Q209L^ using *Plp1-creERT* with tamoxifen at 5 weeks of age and studied the effects of knocking out one copy of *Nf1*. We found that *Nf1* heterozygous loss accelerated the development of intra-dermal and uveal melanoma. We investigated the transcriptional changes that accompany *Nf1* loss in the context of intra-dermal and uveal melanoma and found evidence for up-regulation of cAMP signaling and down-regulation of myogenesis gene expression, respectively. Lastly, we discovered in the process that GNAQ^Q209L^ expression in *Plp1*-expressing cells can drive the formation of neoplasms similar to peripheral nerve sheath tumors, even without *Nf1* loss.

## RESULTS

### Review of 17q11.2 copy number changes in intra-dermal melanocytic lesions

We surveyed the literature for studies that molecularly examined intra-dermal nevi and melanomas, which are commonly called blue-nevus type lesions. Few studies had sequenced the *NF1* locus in these types of lesions. One case, a cellular blue nevus, was mutant for *NF1* (NF1^S856R^) [24]. There were, however, 115 cases with genome-wide copy number analysis (**Table 1**) [25,26,27,28,29,30,31]. These cases were described as benign, intermediate/ambiguous, or malignant in their corresponding publications (color coded in Table 1). Copy number was assessed using array CGH, molecular inversion probe technology, or next generation sequencing. Copy number alterations were absent in the 86 benign or intermediate cases. Four of the 29 malignant cases (14%) exhibited partial loss of chromosome 17, which included 17q11.2 [25,26,27]. This suggested that haploinsufficiency of *NF1* could have a role in the progression of blue nevus type lesions.

**Table One.** 115 intra-dermal melanocytic lesions with available 17q11.2 copy number status. Yellow cell indicates benign lesions, orange cells indicate intermediate grade lesions, and red cells indicate malignant melanoma. ^a^dim(17)(q11-q22), ^b^dim(17)(q11-q22), ^c^dim(17)(pter-q21), ^d^dim(17)(q11.2-q21.31).

| Paper | Year | Method | Case description | Number of cases with 17q11.2 loss | Total number of cases | % |
| --- | --- | --- | --- | --- | --- | --- |
| Maize et al | 2005 | Array CGH | Cellular blue nevus | 0 | 11 | 0 |
| Chan et al | 2016 | Molecular inversion probe | Cellular blue nevus | 0 | 5 |  |
| Costa et al | 2016 | Array CGH | Cellular blue nevus | 0 | 24 |  |
| Held et al | 2013 | Array CGH | Cellular blue nevus with features of deep penetrating nevus | 0 | 15 | 0 |
| Maize et al | 2005 | Array CGH | Atypical blue nevus | 0 | 11 |  |
| Chan et al | 2016 | Molecular inversion probe | Atypical cellular blue nevus | 0 | 12 |  |
| Held et al | 2013 | Array CGH | Ambiguous cellular blue nevus | 0 | 8 |  |
| Goto et al | 2019 | Array CGH | Blue nevus like melanoma mimicking PEM | 0 | 1 | 14% |
| Costa et al | 2016 | Array CGH | Melanoma ex blue nevus | 2<sup>a,b</sup> | 11 |  |
| Shain et al | 2015 | Next gen CNA | Melanoma ex blue nevus | 0 | 1 |  |
| Griewank et al | 2018 | Next gen CNA | Melanoma ex blue nevus | 0 | 2 |  |
| Maize et al | 2005 | Array CGH | Malignant blue nevus/mimic | 1<sup>c</sup> | 5 |  |
| Chan et al | 2016 | Molecular inversion probe | Melanoma ex blue nevus | 1<sup>d</sup> | 9 |  |

### Review of 17q11.2 copy number changes in the TCGA uveal melanoma dataset

We next examined *NF1* in uveal melanoma. Uveal melanoma arises from melanocytes located in the uveal tract of the eye [32]. There are 80 cases of primary uveal melanoma in The Cancer Genome Atlas (TCGA) UVM dataset (https://portal.gdc.cancer.gov) [33]. We examined these cases for *NF1* mutation status and copy number alterations. No somatic point mutations in *NF1* were found in uveal melanoma. However, two cases carried a heterozygous, partial loss of chromosome 17 that included the *NF1* gene. The copy number loss spanned 17q11.2-17q25.2 (*TWF1P1* to *JMJD6*) in case TCGA-VD-AA8M and 17p13.3-q21.32 (*DOC2B* to *HOXB2*) in case TCGA-VD-AA8Q. Both cases carried a typical glutamine substitution at 209 in *GNAQ*. Strikingly, TCGA-VD-AA8Q carried a *RASA2* mutation (Rasa2^K81Q^). This alteration was rated with a SIFT impact score of 0.03 (deleterious) and a Polyphen score of 0.977 (probably damaging). Recurrent mutations at neighboring residue S82 have been previously reported in sun exposed melanomas [5]. The K81Q mutation was the only *RASA2* mutation present in the TCGA-UVM dataset, so its co-occurrence with *NF1* copy number loss is quite interesting, given the previously observed co-occurrence of *RASA2* and *NF1* mutations in cutaneous melanomas [12]. TCGA-VD-AA8Q was reported to end in death by metastatic disease. Both cases lacked mutations in the common UM tumor suppressors, *SF3B1*, *BAP1* and *EIF1AX*. We note also that there are six other cases with copy number gain (3x or 4x) that included the *NF1* locus, with unknown significance.

### Two dermal tumor types found in *GNAQ^Q209L^* expressing mice

We next induced heterozygous loss of *Nf1* in our GNAQ^Q209L^ expressing mouse model as an experimental system to test the role of *Nf1* in non-epithelial melanoma. To drive oncogenic GNAQ^Q209L^ expression in mice, we used the previously described, *Rosa26-floxed stop-GNAQ^Q209L^*(“*R26-fs-GNAQ^Q209L^*”) allele [34]. In this allele, constitutively active human *GNAQ^Q209L^*was knocked into the ubiquitously expressed *Rosa26* locus, preceded by a *loxP* flanked stop cassette that prevents transcription. In cells that express Cre recombinase, the two *loxP* sites are recombined and the intervening stop cassette is deleted, allowing GNAQ^Q209L^ to be expressed. To knockout one copy of *Nf1*, we used the conditional *Nf1^tm1Par^* (“*Nf1^flox^*”) mice [35]. These alleles were combined with the widely used tamoxifen inducible *Plp1-creERT* transgene (*Tg(Plp1-cre/ERT)3Pop)*. This line is expressed in melanocytes and peripheral nerve sheath cells (Schwann cells) and was of interest to us due to its previous connections with *Nf1* and tumorigenesis [22,36,37,38,39,40]. All mice were on an inbred C3HeB/FeJ genetic background.

We induced CreERT activity 5 weeks after birth using twice daily IP injections of tamoxifen for three days. We injected *Plp1-creERT* mice that carried just *R26-fs-GNAQ^Q209L^*/+ (n=14) or *R26-fs-GNAQ^Q209L^*/+ and *Nf1^flox^/+* (n=11), in two sequential cohorts including both males and females. Five *Plp1-creERT*/+; *Nf1^flox^/+* control mice were also injected for comparison. In addition, there were uninjected control mice housed in tamoxifen-free cages: *Plp1-creERT*/+; *R26-fs-GNAQ^Q209L^*/+; *Nf1^flox^/+* (n=2), *Plp1-creERT*/+; *R26-fs-GNAQ^Q209L^*/+ (n=5), and 35 of their cagemates of various other genotypes. The longest surviving mouse among those expressing GNAQ^Q209L^ was 72 weeks old, and therefore all other mice were aged to at least 72 weeks. Nine of the control mice (out of the 47 total) had to be euthanized between 50-63 weeks due to various problems (weight loss mostly, but also head tilt, eye infection, or skin infection), but not for tumors.

The first noticeable phenotype in the mice expressing GNAQ^Q209L^ was hyper-pigmented tail, ear and foot skin, which developed within 8 weeks of the tamoxifen injection (**Figure 1A**). *Nf1* loss greatly increased dermal hyper-pigmentation (**Figure 1B-D**). Over time, as we aged the mice, we found that the *Plp1-creERT*/+; *R26-fs-GNAQ^Q209L^*/+; *Nf1^flox^/+* mice lost weight more quickly than *Plp1-creERT*/+; *R26-fs-GNAQ^Q209L^*/+; +/+ mice (**Figure 1E**). Large tumors (>0.5 cm in diameter) were the primary reason for euthanasia in about half of the mice of both genotypes. The rest were euthanized due to other ill health indicators (piloerection, hunching, excessive scratching, and/or having severely thickened ears), without finding a large tumor.

**Figure 1.**
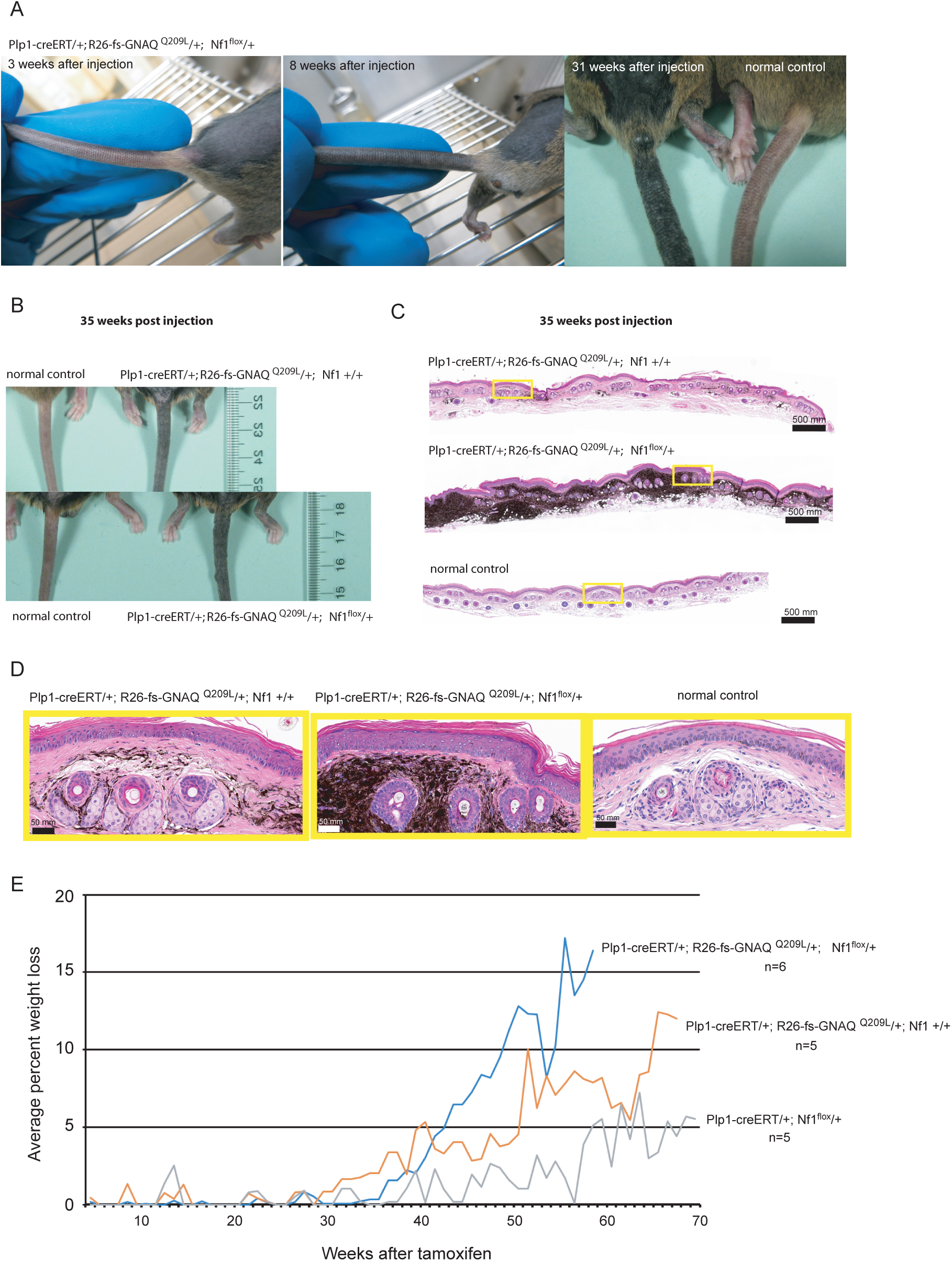
Effects of heterozygous *Nf1* loss on the skin pigmentation and weight of mice expressing oncogenic GNAQ. **(A)** The tail of a *Plp1-creERT*/+; *R26-fs-GNAQ^Q209L^*/+; *Nf1^flox^*/+ mouse injected with tamoxifen at 5 weeks of age, at various indicated time points after injection, showing progressive hyper-pigmentation. (**B**) *Plp1-creERT*/+; *R26-fs-GNAQ^Q209L^*/+; *Nf1^flox^*/+ and *Plp1-creERT*/+; *R26-fs-GNAQ^Q209L^*/+; +/+ mouse tails at 35 weeks post injection, compared to their uninjected littermates. (**C-D**) H&E analysis of tail skin from *Plp1-creERT*/+; *R26-fs-GNAQ^Q209L^*/+; *Nf1^flox^*/+ and *Plp1-creERT*/+; *R26-fs-GNAQ^Q209L^*/+; +/+ mice at 35 weeks following injection, with an uninjected control mouse tail skin section. Yellow boxes in C are enlarged in D. (**E**) A graph of weight loss over time in mice of the indicated genotypes, beginning at tamoxifen injection and continuing until all mice of the genotype had to be euthanized.

The dermal tumors in mice expressing GNAQ^Q209L^ occurred in two distinct types (**Figure 2**). The first type (n=12 tumors) consisted of tumors that lost their overlying fur as they grew and were very darkly pigmented throughout (hereafter referred to as intra-dermal melanomas) (**Figure 2B,C)**. These tumors were S100b positive, which is normal for melanomas (**Supplementary Figure 1A,C**). They also had many CD68-positive macrophages/melanophages (**Supplementary Figure 1B,D**). These results are identical to what we have previously observed when expressing GNAQ^Q209L^ in just melanocytes and aging the mice [39,41].

**Figure 2.**
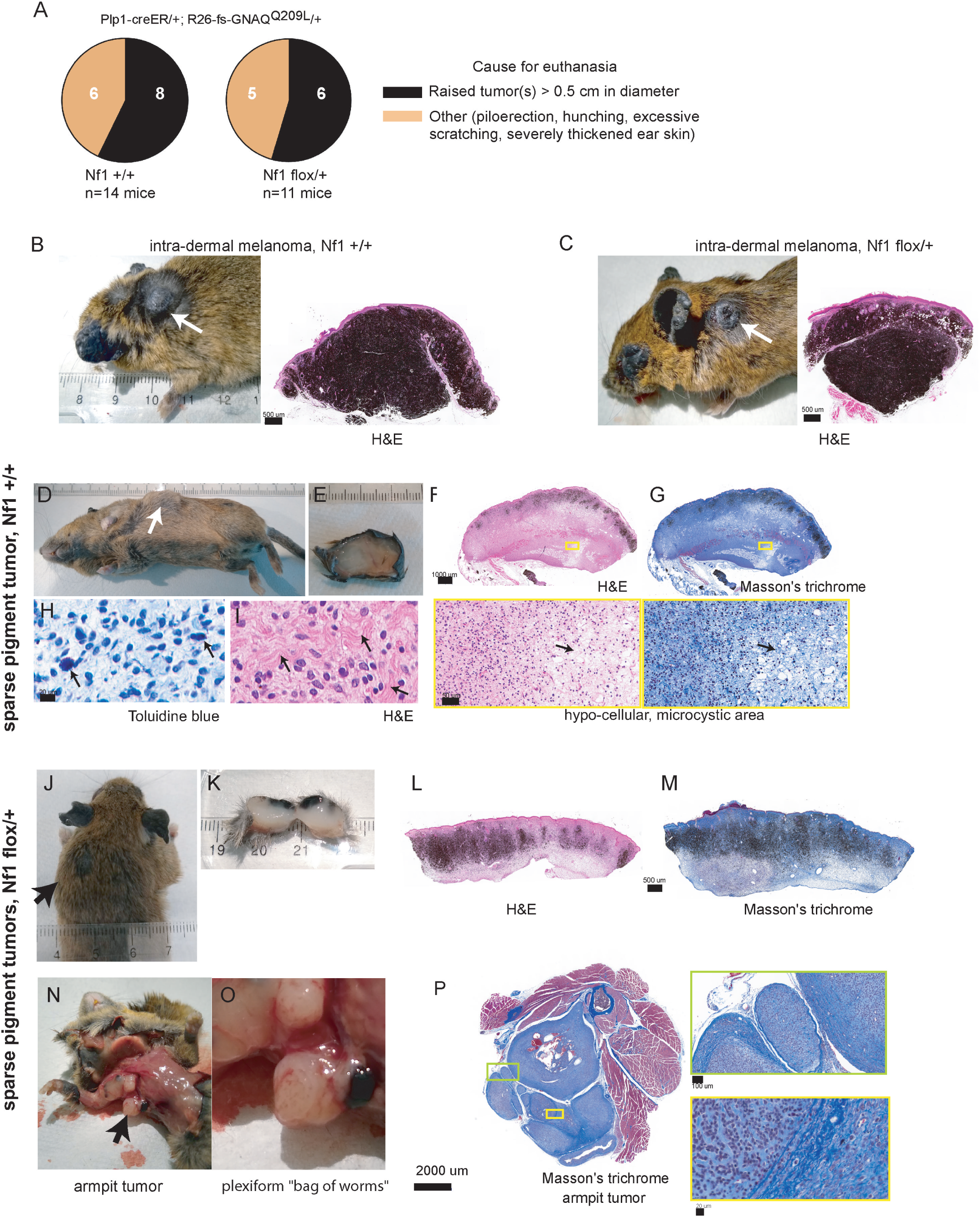
Large tumor formation in mice expressing oncogenic GNAQ using *Plp1-creERT*. **(A)** Pie charts summarizing tumor and non-tumor causes of death of mice in the study, with or without *Nf1* +/− loss. (**B**) 66 weeks after tamoxifen *Plp1-creERT*/+; *R26-fs-GNAQ^Q209L^*/+; +/+ mouse (left) with an intra-dermal melanoma on the neck. The tumor is shown stained with H&E (right). (**C**) 51 weeks after tamoxifen *Plp1-creERT*/+; *R26-fs-GNAQ^Q209L^*/+; *Nf1^flox^*/+ mouse (left) with an intra-dermal melanoma on the upper back. The tumor is shown stained with H&E (right). (**D-G**) 61 weeks after tamoxifen *Plp1-creERT*/+; *R26-fs-GNAQ^Q209L^*/+; +/+ mouse (D) with a sparse pigment tumor on the side of the trunk; tumor shown just after removal (E), and sectioned and stained with H&E (F) and Masson’s trichrome (G). An area containing both hypo and hyper cellular regions of the tumor is enlarged in the boxes below, arrows. (**H**) Close up images of granular mast cells (arrows) in toluidine blue stained section of a sparse pigment tumor. (**I**) Wavy collagen bundles (arrows) in H&E stained section of sparse pigment tumor. (**J**) 48 weeks after tamoxifen *Plp1-creERT*/+; *R26-fs-GNAQ^Q209L^*/+; *Nf1^flox^*/+ mouse that developed a sparse pigment tumor on the left shoulder (arrow). (**K**) Another sparse pigment tumor from a different *Plp1-creERT*/+; *R26-fs-GNAQ^Q209L^*/+; *Nf1^flox^*/+ mouse, cut in half. (**L,M**) The tumor in J was sectioned and stained with H&E (L) and Masson’s trichrome (M). (**N,O**) The mouse in J also had an armpit tumor (N). This tumor had nodules with a macroscopic “bag of worms” plexiform appearance (O). (**P**) Masson’s trichrome staining of the nodule shown in O, with enlargements of areas of interest. The surrounding muscle and bone are included.

The second intra-dermal tumor type has not been seen before in mice, to our knowledge. These tumors were broader and flatter in shape and did not disrupt the overlying fur (**Figure 2D, J).** They also contained much less pigment, hence they are referred to hereafter as “sparse pigment” tumors. When dissected and cut in half, sparse pigment tumors were grey colored and glistened, and had a more rubbery texture than the intra-dermal melanomas (**Figure 2E, K**). The sparse pigment tumors were encapsulated and had alternating areas of increased and decreased cellularity (**Figure 2F, G, L, M**). Toluidine blue staining revealed granular mast cells (**Figure 2H**), while H&E staining and Masson’s trichrome staining showed abundant wavy collagen filaments (**Figure 2I**). Blood vessels were more hyalinized in the sparse pigment tumors than the intra-dermal melanomas (**Supplementary Figure 2**). No whorls or verocay bodies were observed.

In one interesting *Plp1-creERT*/+; *R26-fs-GNAQ^Q209L^*/+; *Nf1^flox^/+* mouse, two sparse pigment tumors developed in the dermis, one much larger than the other (**Figure 2J**). Around the same time, the mouse starting holding up its right foreleg. It was euthanized and necroscopy revealed a tumor in the right armpit as well (**Figure 2N**). The armpit tumor was composed of three connected nodules, two of which had a plexiform “bag of worms” macroscopic appearance that is characteristic of plexiform neurofibromas (**Figure 2O**). The Masson’s trichrome staining in **Figure 2P** shows a possible connection of the nodule in Fig. 2O to the body, as well as some of the tumor lobes and structures making up the plexiform macroscopic appearance. Our working hypothesis is that these sparse pigment tumors arise from Schwann cells, and they are related to neurofibromas, schwannomas or another kind of peripheral nerve sheath tumor. Altogether, we observed 8 sparse pigment tumors.

We noted a trend in the locations of the intra-dermal melanomas versus sparse pigment tumors. The intra-dermal melanomas were most frequently found around the back of the neck, while the sparse pigment tumors were more frequently found on the sides of the trunk (**Figure 3A**). The ratio of intra-dermal melanomas to sparse pigment tumors was similar in *Nf1* +/+ versus *Nf1^flox^*/+ mice (**Figure 3B**) and both types were found in males and females (**Supplementary Table 1a**). Kaplan-Meier analysis showed that *Nf1^flox^/+* significantly decreased tumor free survival (p = 0.0016; **Figure 3C, Supplementary Table 1a**).

**Figure 3.**
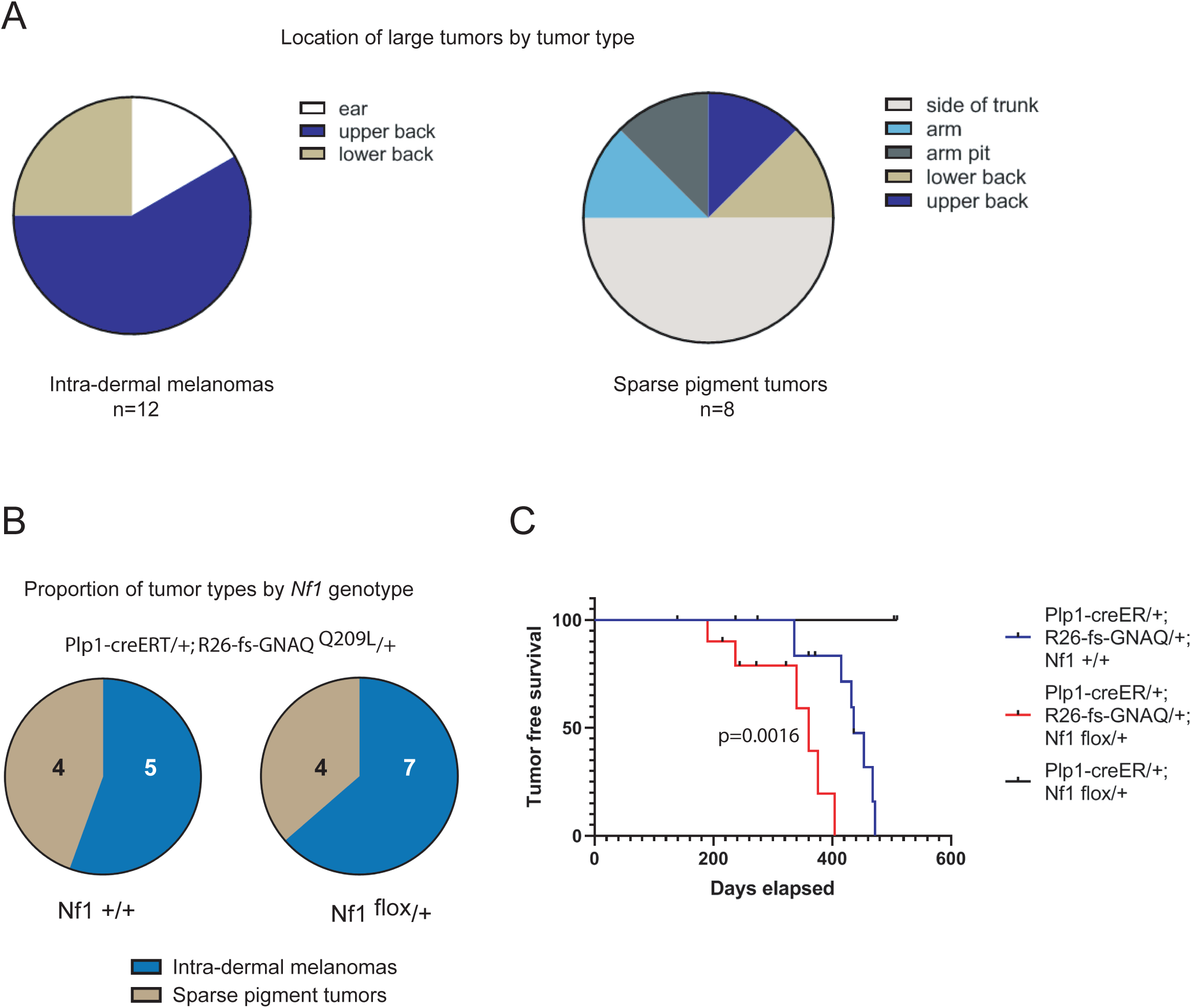
Tumorigenesis is accelerated by *Nf1* +/− loss in *Plp1-creERT/+*; *R26-fs-GNAQ^Q209L^/+* mice. **A)** Location of large intra-dermal tumors on the body, by tumor type (intra-dermal melanoma or sparse pigment). **B)** Proportion of tumors of each type, by *Nf1* genotype. **C)** Kaplan Meier plot of tumor free survival in *Plp1-creERT*/+; *R26-fs-GNAQ^Q209L^*/+; *Nf1^flox^*/+ mice versus *Plp1-creERT*/+; *R26-fs-GNAQ^Q209L^*/+; +/+ mice. There was a significantly decreased tumor free survival in *Plp1-creERT*/+; *R26-fs-GNAQ^Q209L^*/+; *Nf1^flox^*/+ mice. For individual mouse data, see Supplementary Table 1a.

### *Nf1* loss accelerated tumorigenesis in the mouse eye

To assess uveal melanoma in the mice, we collected eyes and stained sections taken at the middle of the eyes with H&E. In all mice expressing GNAQ^Q209L^, there was an expansion of the pigmented tract (**Figure 4A**). We bleached sections to remove the melanin and performed IHC for S100b (for melanocytes) or RPE65 (for the retinal pigment epithelium) to confirm that the excess pigmented tissue was from the uveal tract. RPE65 stained an expected narrow strip of cells between the neural retina and the choroid (**Figure 4C**). The tumor tissues were positive for S100b and negative for RPE65, therefore, uveal melanoma. The *Plp1-creERT/+; Nf1^flox^/+* control mice injected with tamoxifen exhibited normal eyes (**Figure 4A**).

**Figure 4.**
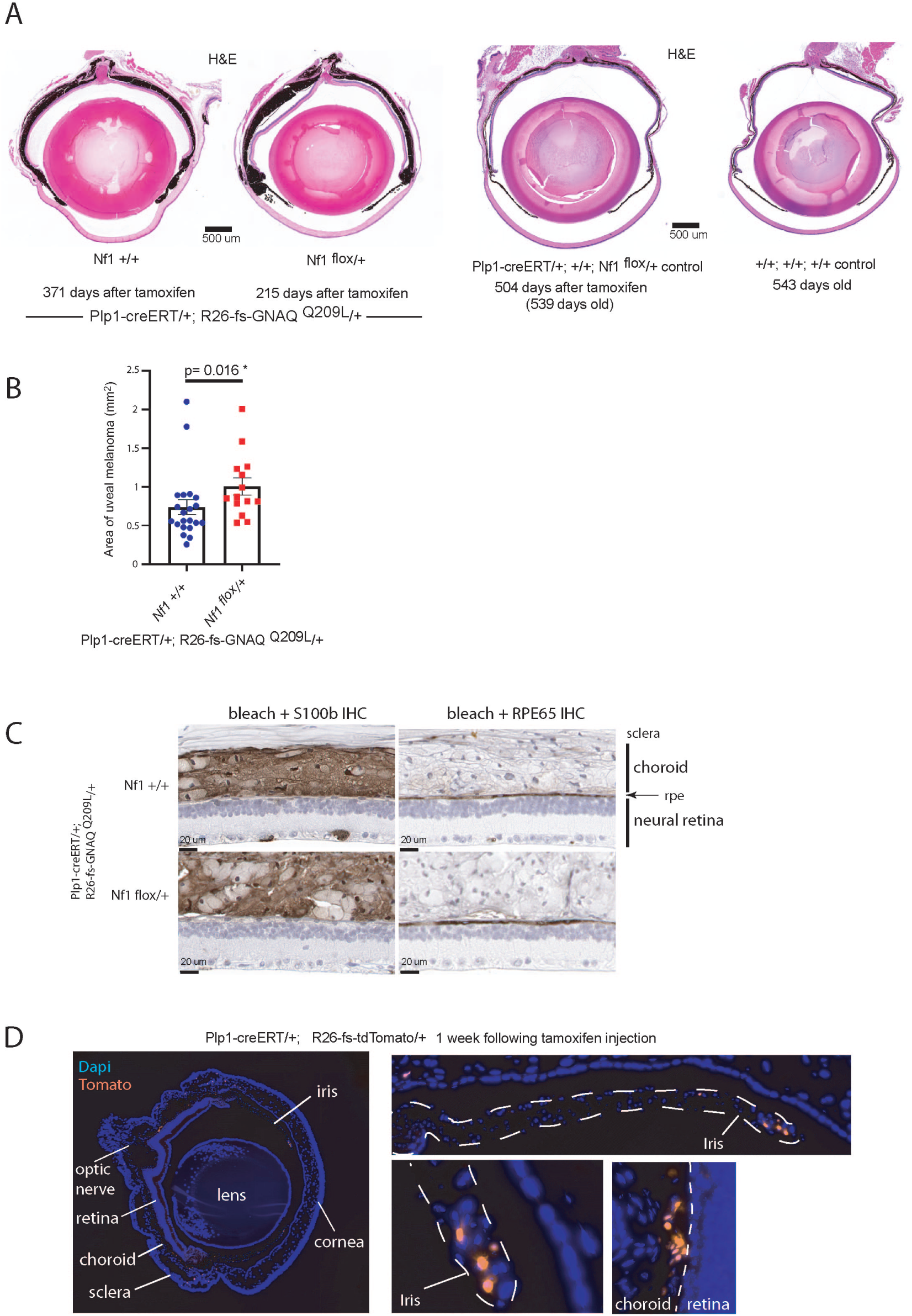
Uveal melanomagenesis is accelerated by *Nf1* +/− loss in *Plp1-creERT/+*; *R26-fs-GNAQ^Q209L^/+* mice. **A)** H&E stained sections of representative eyes of the indicated genotypes. **B)** Quantification of the area of pigmented tissue in *Plp1-creERT*/+; *R26-fs-GNAQ^Q209L^*/+; *Nf1^flox^*/+ versus *Plp1-creERT*/+; *R26-fs-GNAQ^Q209L^*/+; *Nf1* +/+ eyes. There was a significant increase in area in the *Nf1^flox^/+* eyes. **C**) IHC to determine whether the excess pigmented tissue in eyes was uveal melanoma, using IHC to examine expression of S100B (for melanocytes, left) or RPE65 (for retinal pigment epithelium ‘RPE’, right). The excess tissue was S100B-positive and the RPE was a normal, single layered epithelium. **D**) 6 week *Plp1-creERT/+; Rosa26-fs-tdTomato/+* eye 1 week following injection with tamoxifen. *Plp1-creERT* induced tomato expression (red) in cells in the uveal tract, including the iris, ciliary body and choroid (shown in enlargements, right). Sections were counterstained with DAPI (blue).

The size of the uveal melanomas was quantified using ImageJ. There was a significant increase in the average size of the uveal melanoma in the *Nf1^flox^*/+ eyes compared to the *Nf1* +/+ eyes (p = 0.016, **Figure 4B**). Note that these eyes were collected at the humane or dermal tumor endpoints described in the previous section, not at a fixed time point. The average number of days of survival after tamoxifen in the *Nf1^flox^*/+ mice whose eyes were sectioned was 98 days less than the *Nf1* +/+ mice, yet the uveal melanomas were bigger on average in *Nf1^flox^*/+ mice.

We crossed *Plp1-creERT* to the *R26-fs-tdTomato* reporter line [*Gt(ROSA)26Sor^tm14(CAG-tdTomato)Hze^*] to label cells expressing *Plp1-creERT* in the eye. We administered tamoxifen by IP injection at 5 weeks and collected the eyes at 6 weeks. There were tdTomato positive cells in the iris and choroid of the eyes, but not in the sclera or retina (**Figure 4D**). Hence, *Plp1-creERT* can be used to induce uveal melanoma when tamoxifen is given at 5 weeks of age. Furthermore, *Nf1* haploinsufficiency accelerated uveal melanomagenesis driven by GNAQ^Q209L^.

### Other pigmented lesions in the mice

Oncogenic GNAQ also causes a variety of other pigment cell defects. GNAQ is frequently mutant in primary melanoma of the central nervous system (CNS) [8,39]. To assess the pigmented lesions associated with the CNS in the mice, we photographed the spines and collected the brains during necroscopy of the second cohort of the *Plp1-creERT/+; R26-fs-GNAQ^Q209L^*/+ mice, with and without conditional *Nf1/+* loss. All mice expressing GNAQ^Q209L^ exhibited small pigmented lesions in the lumbar area of the back, centered above the spine (**Supplementary Figure 3A,B**). There were also multiple examples of mice with pigmented lesions within the spine upon sectioning (**Supplementary Figure 3C**). However, there was no significant difference between *Nf1^flox^/+* and +/+ mice. In addition, we noted lesions on the ventral brain surface, but again there was no significant difference between *Nf1^flox^/+* and +/+ mice with respect to lesion size (**Supplementary Figure 3D, E**). Compared to the *Mitf-cre*/+; *R26-fs-GNAQ^Q209L^*/+ model described previously [34,41], the CNS lesions were much smaller and there also was no hyperactivity or head tossing.

GNAQ^Q209L^ expressing mice also develop pigmented lung lesions. It is difficult to know whether these are distant metastases, but there are no known resident melanocytes in the lungs [34]. In animals expressing GNAQ^Q209L^, there were on average 19 lung lesions per mouse, regardless of the *Nf1* genotype (**Supplementary Figure 4**). There was no difference in the average size of the lung lesions in the *Nf1* +/+ versus *Nf1^flox^/+* mice. These lesions were all small, less than 0.13 mm^2^ in area (**Supplementary Figure 4B-D**).

Lastly, we examined the phenotype of uninjected mice of the mutant genotypes. We previously reported that there is some limited activation of CreERT in *Plp1-creERT/+; R26-fs-GNAQ^Q209L^*/+ mice in the absence of any tamoxifen (*i.e.* “leaky CreERT activity”) [39]. This small level of activity can be detected because changes in pigmentation by GNAQ^Q209L^ are easy to spot. Here, we examined two *Plp1-creERT*/+; *R26-fs-GNAQ^Q209L^*/+; *Nf1^flox^/+* mice and five *Plp1-creERT*/+; *R26-fs-GNAQ^Q209L^*/+; *+/+* mice, housed in tamoxifen-free cages, after aging to 72 weeks. As in [39], there were tiny punctate spots on all shaved trunks. In addition, two tails exhibited small pigmented lesions in the dermis (**Supplementary Figure 5**), one eye had an expanded pigmented layer (**Supplementary Figure 6**), and one spine had an overlying lesion within the muscle (**Supplementary Figure 7**).

### Effects of *Nf1* loss on the melanoma transcriptome

We next performed bulk RNAseq to identify differentially expressed (DE) genes caused by *Nf1* haploinsufficiency. We included 7 intra-dermal melanomas (3 *Nf1* +/+ and 4 *Nf1^flox^*/+), 4 sparse pigment tumors (all *Nf1^flox^*/+), and 10 uveal melanomas (5 *Nf1* +/+ and 5 *Nf1^flox^*/+). RNAseq was performed on all 21 RNA samples in one run. We used DESeq2 to compare the samples in several different analyses, described below. To assess overall relationships, we included all samples for unsupervised clustering by gene expression, regardless of the *Nf1* genotype or tumor type. In the principal component analysis (PCA) (**Figure 5A**) and sample distance dendrogram (**Figure 5B**), the three tumor types clustered separately. We included the plexiform armpit tumor and a sparse pigment tumor from the same mouse for RNAseq (the mouse shown in **Figure 2J,N**). The armpit tumor grouped with the other sparse pigment tumors in gene expression, despite its different location in the body (indicated by asterisks in Figure 5B). The *Nf1* genotypes are indicated by color in the same PCA plot in **Figure 5C**, which revealed no strong pattern of clustering by genotype, compared to the big differences produced by tumor type.

**Figure 5.**
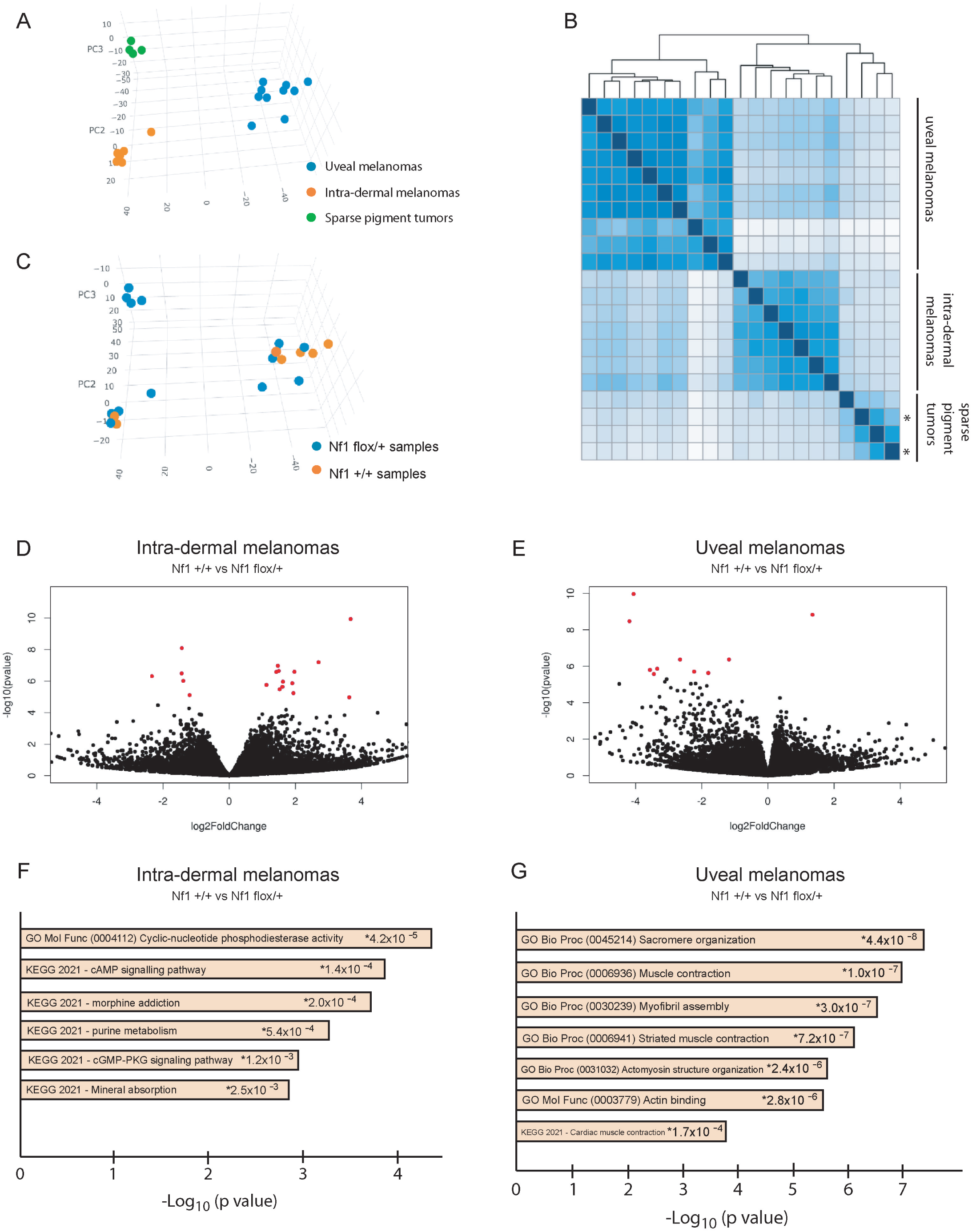
Transcriptomic analysis of uveal melanoma, intra-dermal melanoma, and sparse pigment tumors. **(A,B)** All samples were included in unsupervised clustering by gene expression, regardless of the *Nf1* genotype or tumor type. Principal component analysis (A) and the sample distance dendrogram (B) shows that the three different tumor types cluster separately from each other. In B, the two sparse pigment tumors (shown in Figure 2J and 2N) from the same mouse are indicated with asterisks. **C)** Same PCA plot as shown in A, but now colored coded to show the *Nf1* genotypes. There was no obvious clustering by *Nf1* genotype within tumor types. **D,E)** Volcano plots of differential gene expression in *Nf1^flox^*/+ versus *Nf1 +/+* intra-dermal melanoma (D) and uveal melanomas (E). **F,G)** Top terms found in GO analysis of the differentially expressed genes in intra-dermal melanoma (F) and uveal melanoma (G).

We then performed DE analysis to compare *Nf1^flox^*/+ versus *Nf1 +/+* tumors within tumor types. Volcano plots of the results are shown in **Figures 5D** (intra-dermal melanoma) and **Figure 5E** (uveal melanomas). In the intra-dermal melanomas, there were 15 up-regulated genes and 10 down-regulated genes in the *Nf1^flox^/*+ tumors compared to *Nf1* +/+, with p adj < 0.05 (**Supplementary Table 1b**). Among these genes, the most significant term was GO (gene ontology) molecular function:0004112, ‘Cyclic nucleotide phosphodiesterase activity’, p=4.2×10^−5^ (**Figure 5F, Supplementary Table 1c**). This was supported by the up-regulation of *Adcy1* and *Atp1b2* and the down-regulation of *Pde10a* and *Pde3a* in the *Nf1^flox^*/+ tumors. cAMP levels are regulated by adenylyl cyclases (Adcy’s), which produce cAMP from ATP, and by the activity of phosphodiesterases (PDEs), which hydrolyze and degrade cyclic nucleotides (cAMP and cGMP). The balance between the activity of these two enzymatic families controls the activation of the downstream effectors, protein kinase A and the cAMP responsive element binding protein (CREB).

In the uveal melanomas, there were 23 down-regulated genes and 2 up-regulated DE genes with p_adj_ < 0.05 (**Supplementary Table 1d**). There was an exceptionally clear signal from the *Nf1* mutant uveal melanomas in GO analysis (**Figure 5G, Supplementary Table 1e**). 20 of the 23 down-regulated genes were related to muscle contraction and/or myogenesis. The most significant term was GO biological process:0045214, ‘Sacromere organization’, p = 4.4×10^−8^. The complete list of DE genes relating to muscle function were: *Acta1*, *Actn3*, *Atp2a1*, *Casq1*, *Ckm*, *Cmya5*, *Cox6a2*, *Dhrs7c*, *Mybpc2*, *Myh4*, *Mylk2*, *Myom2*, *Myoz1*, *Pgam2*, *Phkg1*, *Pvalb*, *Synpo2*, *Tnni2*, *Tnnt3*, and *Tpm2*. The relevance of cAMP and muscle gene expression changes can be found in the discussion section.

Although UM prognosis is very strongly correlated with monosomy 3 and the loss of *BAP1*, we wondered whether any of the DE genes identified above were correlated with survival in the TCGA-UVM database. In fact, 20% of these DE genes (9/49) were significantly correlated with UM survival, suggesting there could be some overlap in prognosis related targets by different genetic pathways (**Figure 6**). For all but one gene, the relationship was in the expected direction (for example, if the gene was up-regulated in the *Nf1^flox^/+* mouse tumors, then higher expression was correlated with a worse outcome in human patients). Two highly significant correlations were for the muscle *COX6A2* gene (p=1.7×10^−5^) and for *ADCY1* (p=7.5×10^−5^). Also, a very significant correlation was found for the up-regulated *GFRA2* gene (p=5.9×10^−6^). Expression of *GRFA2* designates a neural crest stem cell signature found in common cutaneous melanomas [42]. There was only one DE gene that was found in both tumor types when *Nf1* was mutant. This was *Rpl26*, which encodes a large subunit ribosomal protein. Although RPL26 has been previously linked to cancer through the regulation of p53, the prediction is that *Rpl26* down-regulation would promote tumorigenesis and *Rpl26* was up-regulated in our datasets [43].

**Figure 6.**
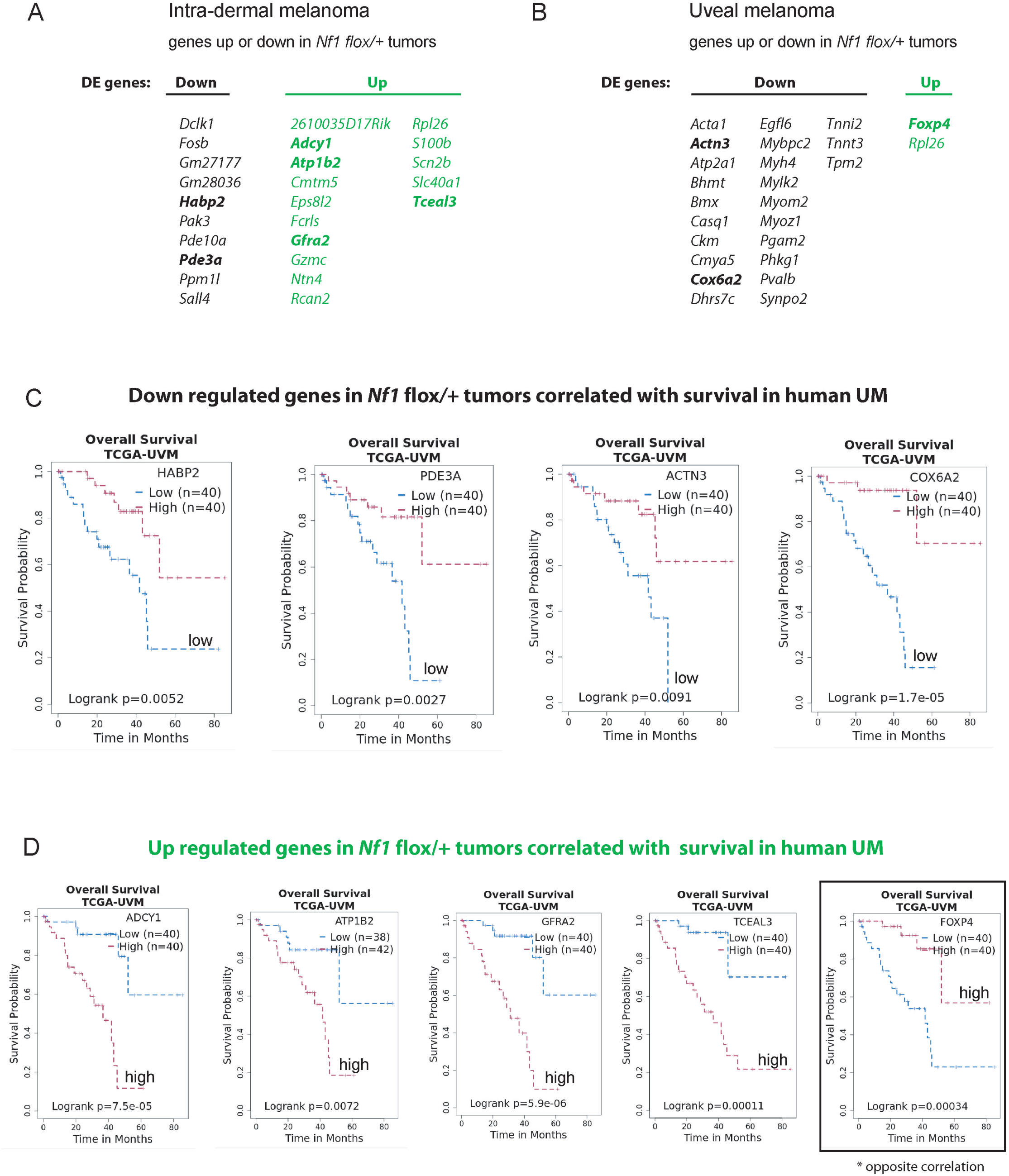
The expression of differentially expressed genes in *Nf1^flox^*/+ melanomas is associated with survival in human uveal melanoma (TCGA-UVM). **A,B)** List of all significant up- and down-regulated genes in *Nf1^flox^*/+ intra-dermal (A) or uveal (B) melanoma (p_adj_ < 0.05). Bolded genes are associated with survival in human uveal melanoma. **C,D)** Kaplan-Meier survival plots of down-regulated (C) and up-regulated (D) differentially expressed genes that were significantly associated with survival in the TCGA-UVM dataset. Genes whose low expression was associated with worse survival were down-regulated in *Nf1^flox^*/+ tumors, and genes whose high expression was associated with worse survival were up-regulated in *Nf1^flox^*/+ tumors, except for one gene, *Foxp4*.

### Transcriptomic differences between intra-dermal melanoma and sparse pigment tumors

We then compared gene expression in the intra-dermal melanomas and sparse pigment tumors, irrespective of *Nf1* genotype, in order to better define their differences. There were 7350 DE genes with a p_adj_ cut off < 0.05 (**Supplementary Table 1f**). We first considered the top genes that were most differentially expressed by log_2_fold change. Many well known pigmentation genes were among the 150 most up-regulated genes in the intra-dermal melanomas: *Slc45a2, Tyrp1, Tyr, Pmel, Dct, Oca2, Mlana, Slc24a4*, and *Mc1r*. On the other hand, many Schwann cell/oligodendrocyte/glia supporting genes were in the top 150 gene up-regulated in the sparse pigment tumors: *Gpr17, Crispld1, Ptprz1, Col20a1, Scn7a, Lrrn1, Wnt16, Matn4, Asic4, Mog, Nkx2-2, Kirrel3, Tenm3, Kcnh8, Dbh, Srcin1*, and *Plxnb3* (**Supplementary Table 1g).** We also looked up various genes classically expressed by Schwann cell precursors or Schwann cells and found that *Gap43*, *Fabp7*, *Mpz*, *Dhh*, *Ngfr*, *Ncam1*, and *Mbp* were all significantly up-regulated in the sparse pigment tumors compared to the intra-dermal melanomas, as was *Plp1* (*Proteolipid protein 1*). The top 150 up-regulated genes in the sparse pigment tumors returned the significantly enriched terms, ‘Schwann cells in adrenal,’ ‘Oligodendrocytes in cerebrum’, ‘Schwann cells in muscle,’ ‘Oligodendrocytes in cerebellum’, *etc* (**Figure 7A**). This supports the hypothesis that the sparse pigment tumors are some kind of nerve sheath neoplasm.

**Figure 7.**
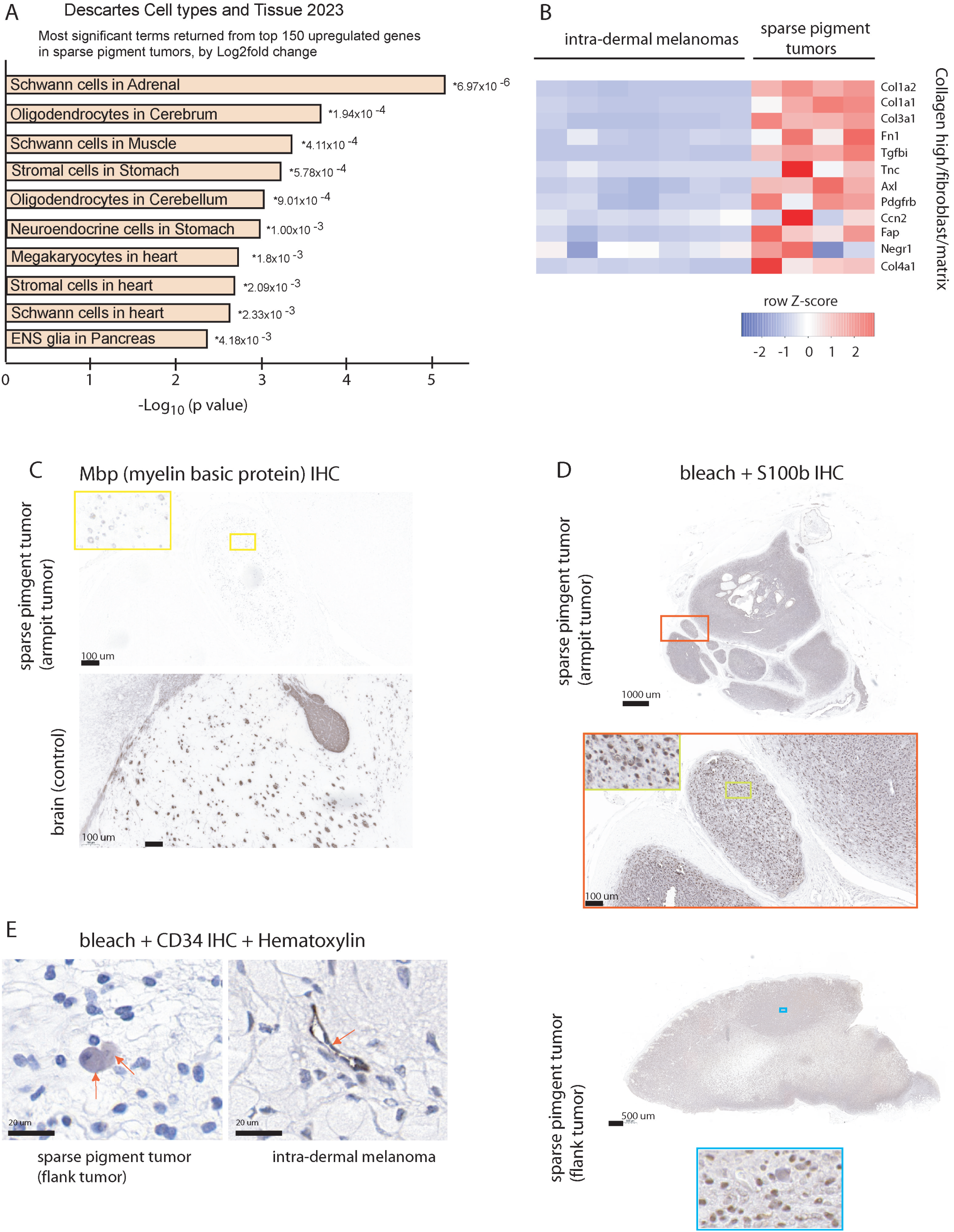
Gene expression analysis suggests that sparse pigment tumors are related to schwann cells. **A)** The most significant terms returned from GO analysis of the top 150 genes up-regulated in sparse pigment tumors versus intra-dermal melanomas, by log2fold change, using Descartes cell types and tissues 2023. **B)** Heat map of gene expression that indicates higher collagen, fibroblast, and matrix gene expression in sparse pigment tumors. **C,D,E)** IHC for myelin basic protein (Mbp), a marker for Schwann cells (C), or S100b, a marker for neural crest derived cells, (D), or CD34 (E), in indicated tissues. Red arrows in E indicate cells of interest. The CD34 IHC gave different results in the two tumor types. CD34 positive cells in the intra-dermal melanoma line capillaries and vessels. In the sparse pigment tumors, arrows indicate examples of weak CD34-positive cells that are also round and basophilic.

Miskolczi *et al* found that the presence of fibroblast tumour growth factor-β suppressed YAP/PAX3-mediated MITF expression and was associated with a de-differentiated phenotype in common cutaneous melanoma [44]. We noticed that the second most significant DE gene in our comparison was *Tgfbi* (*Transforming growth factor, beta induced*), which had a log_2_fold change of 4.9 (up-regulated in sparse pigment tumors) and a p_adj_ value of 7.5 × 10^−86^. *Mitf*, on the other hand, was down regulated in the sparse pigment tumors with a log_2_fold change of −3.15 and a p_adj_ value of 1.23 × 10^−15^. Other genes in the suggested collagen high/fibroblast/matrix gene signature proposed in reference [44] were up-regulated in the sparse pigment tumors compared to the intra-dermal melanomas (genes in heatmap, **Figure 7B**). This could reflect a greater contribution of fibroblasts and extracellular matrix (ECM) in the sparse pigment tumors.

Myelin basic protein (Mbp) is a marker for peripheral glial cells. The only sparse pigment tumor with Mbp-positivity was the armpit tumor (**Figure 7C**). Some of the cells in this tumor expressed Mbp in a ring, such as might be expected in a myelinated nerve. Nerve bundles can become entrapped in plexiform neurofibromas. Mbp expression was weaker than in a positive control normal mouse brain (**Figure 7C**). In contrast, there was widespread and clear S100b staining in all of the sparse pigment tumors (examples shown in **Figure 7D**). S100b expression has been noted as a consistent marker for peripheral nerve sheath tumors. We were also interested in CD34, which has been proposed to distinguish neurofibroma from desmoplastic melanoma by a fingerprint pattern of expression in the neurofibromas [45]. In the mouse intra-dermal melanomas, CD34 positive staining was found in cells lining blood vessels, as mentioned in ref [45] (**Figure 7E**). An endothelial pattern was not clear in the sparse pigment tumors. Instead, there may have been weak CD34 expression in round basophillic cells of unknown identity, which were scattered throughout (**Figure 7E**). In this way, CD34 was differential between the tumor types, but not in a classic neurofibroma pattern.

## DISCUSSION

MAP Kinase (MAPK) pathway activation is one of the key events in melanoma, as well as in many other cancers [1]. Mutations in MAPK pathway components in sun exposed melanoma occur in the oncogenes, *BRAF* and *NRAS*, and the tumor suppressors, *RASA2* and *NF1* [2,3,4,5]. The net intensity of MAPK signaling in a cell is determined by the cumulative activity of components all along the pathway. In sun exposed skin, melanocytic nevi begin with a single activating MAPK mutation, most often in *BRAF*. Multiple and independent genetic alterations incrementally accumulate over time, building to higher levels of MAPK signaling in melanoma [13,46]. This occurs even in the absence of selective pressure by anti-MAPK tumor therapies.

In melanomas that arise outside of an epithelium, such as in the dermis and eye, MAPK pathway activation is achieved through activation of Gα_q/11_ signaling, rather than directly through *BRAF* or *NRAS*. Gα_q/11_ activate phospholipase C-beta, stimulate protein kinase C and feed into the MAPK pathway through RASGRP3 [10]. A tumor promoting effect of neurofibromin loss in non-epithelial melanoma has been suggested by a greater than expected frequency of uveal melanoma in patients with Neurofibromatosis type 1 (*NF1* +/− carriers) [20]. We also previously described a neurofibromatosis type 1 patient who developed a GNAQ^Q209P^-mutant uveal melanoma [18].

To address whether mutations in *NF1* might synergize with Gα_q_ to promote non-epithelial melanomagenesis, we surveyed the published literature and the TCGA uveal melanoma dataset. We found that partial loss of chromosome 17 including the *NF1* gene has been recurrently found in uveal melanoma (2.5%) and blue nevus-type intra-dermal melanoma (14%). The TCGA dataset is a deep source of information and we found that the two cases of uveal melanoma that carried a partial loss of chromosome 17 exhibited a striking array of similarities [33]. Both cases carried a GNAQ^Q209P^ mutation and were wildtype for *BAP1*, *EIF1AX* and *SF3B1*. Each case carried only one other chromosomal copy number abnormality apart from the partial loss of chromosome 17. They were grouped in copy number cluster 1, despite lacking copy number alterations on chromosome 6 that otherwise characterize cluster 1 [33]. In addition, one of the partial loss of chromosome 17 uveal melanoma cases also carried a *RASA2* mutation (Rasa2^K81Q^). Fifty percent of *RASA2* mutant sun-exposed melanomas exhibit a co-occurring mutation in *NF1* [5,12]. This was the only case in the TCGA uveal melanoma dataset with a *RASA2* mutation. Death occurred less than two years after diagnosis. Given this poor outcome and the relative lack of other distinguishing molecular features in these two uveal melanoma cases, the loss of chromosome 17 should be further investigated as a potentially significant finding in uveal melanoma.

It should also be noted that *NF1* is frequently mutated in a rare type of melanoma known as desmoplastic melanoma. Desmoplastic melanoma arises in the elderly on sun exposed skin [47]. This melanoma is primarily located in the dermis, however some cases have an overlying lentigo maligna component. Desmoplastic melanoma is characterized by unpigmented, spindle shaped melanocytes surrounded by an abundant fibrous collagen stroma. Collagen produced by fibroblasts causes an unusual fibrotic appearance, which can be mistaken for a scar. Shain *et al* reported that 54% of desmoplastic melanomas are *NF1* mutant [30,48]. In about half of these cases, *NF1* was heterozygous. *p53* mutations were detected in 48% of cases, with no apparent relationship to *NF1*. Other MAPK hits included *BRAF*, *MAP2K1*, and *MAP3K1,* each in about 5% of cases. *GNAQ* and *GNA11* mutations were not found.

To directly test the interaction between neurofibromin and Gα_q/11_ in a model system, we studied the effects of conditional *Nf1* loss in mice expressing human oncogenic GNAQ^Q209L^ [34]. Because the loss of *NF1* in the human melanoma cases was heterozygous, we generated *Nf1* haploinsufficiency in the mice. We chose to use the *Plp1-creERT* transgene [(*Tg(Plp1-cre/ERT)3Pop)*] for this. This tamoxifen inducible CreERT line is expressed in peripheral glia (Schwann cells) and melanocytes and was of interest to us due to its previous connections with *Nf1* and tumorigenesis [22,36,37,38,39,40]. *Plp1-creERT* has not been used before in post-natal mice to drive *R26-fs-GNAQ^Q209L^*, however, we previously described CNS melanoma with tamoxifen injections early in embryogenesis [39]. Here, we injected tamoxifen at 5 weeks old.

About half of the *Nf1^flox^*/+ and *Nf1*-wildtype mice developed a large tumor on the body (>0.5 cm). The tumors occurred in two types, in both genotypes. One of the types was uniformly and strongly pigmented throughout, and disrupted the growth of the overlying hair. This type is identical to others we’ve observed in GNAQ^Q209L^ expressing mice before and consider intra-dermal melanoma [34,41]. The second type was new. While also intra-dermal, these tumors were deeper, did not disrupt the overlying hair, and had features suggestive of Schwann cell based neoplasms, such as a neurofibroma, schwannoma, or other peripheral nerve sheath tumor. These features were the macroscopic appearance, the combination of dermal and plexiform presentations, alternating areas of hyper- and hypo-cellularity, strong S100b positivity, the presence of mast cells, collagen bundles, and hyalinized vessels, and the up-regulation of Schwann cell specific gene expression compared to the intra-dermal melanomas. These tumors lacked a whorly organization, verocay bodies and CD34 positive fingerprints, which are often found in neurofibromas, so we are not definitive about their exact nature.

In neurofibromatosis type 1, neurofibromas can be dermal/cutaneous or plexiform, but most are of the cutaneous type. Doctors often see patients with a solitary (*i.e.* sporadic) cutaneous neurofibroma in the absence of genetic disease, but multiple cutaneous neurofibromas or a single plexiform neurofibroma strongly suggests a *NF1* +/− carrier status in a patient. *GNAQ* mutations have not been previously reported in either solitary or *NF1*-associated neurofibromas. However, Chen *et al* reported that the activation of Yap in Schwann cells through knockout of *Lats1* and *Lats2* drove cutaneous neurofibroma formation in mice when combined with *Nf1* loss [49]. In melanocytes, Gα_q/11_ activates both Yap and MAPK signaling and therefore GNAQ^Q209L^ could be sufficient to transform Schwann cells [6,34,50,51]. The signaling pathways activated by GNAQ^Q209L^ in Schwann cells still need to be investigated to determine if they are similar to melanocytes, but the known connections between Schwann cells and melanocytes support this. It is also interesting that the one plexiform armpit tumor arose in a mouse that was *Nf1^flox^/+*, but more mice are needed to replicate this finding.

Also of interest are malignant melanotic nerve sheath tumors (MMNSTs), a tumor type redefined in the 2021 WHO classification of tumors of the central nervous system (previously called melanotic schwannoma). These are rare and aggressive neoplasms that frequently have loss of function mutations in the *PRKAR1A* gene, which encodes a regulatory subunit of protein kinase A. Terry *et al* recently published a case report of a woman with MMNST that contained both a *PRKAR1A* frameshift mutation and a GNAQ^R183L^ oncogenic hotspot mutation [52]. Our results support their hypothesis that GNAQ activation can promote tumorigenesis in Schwann cells.

In our experiments, two non-overlapping sets of differentially expressed genes were produced by *Nf1* heterozygous loss in melanoma driven by GNAQ^Q209L^. In the intra-dermal melanomas, the most significant GO term was ‘Cyclic nucleotide phosphodiesterase activity’, p=4.2×10^−5^. This was supported by the up-regulation of *Adcy1* and *Atp1b2* and the down-regulation of *Pde10a* and *Pde3a* in the *Nf1^flox^*/+ tumors. cAMP levels are regulated by adenylyl cyclases (Adcy’s), which produce cAMP from ATP, and by the activity of phosphodiesterases (PDEs), which hydrolyze and degrade cyclic nucleotides (cAMP and cGMP). The balance between the activity of these two enzymatic families controls the activation of the downstream effectors, protein kinase A and the cAMP responsive CREB protein. Higher *ADCY1* expression is very strongly associated with decreased uveal melanoma patient survival (p=7.5×10^−5^). cAMP signaling has not been specifically investigated in non-epithelial melanomas, but otherwise has been the subject of much work in the melanocyte field [53]. In melanocytes, *NF1* inactivation was previously linked to increased activity of cAMP-mediated PKA and ERK, which led to the over-expression of Mitf [16]. Also, malignant peripheral nerve sheath tumor cell lines derived from individuals with neurofibromatosis type 1 were found to have basal cAMP levels two-fold higher than normal Schwann cells [54] and a muscle specific *Nf1* knockout in mice increased cAMP levels [55]. On the other hand, many studies in the brain show the opposite effect, with neurofibromin loss reducing adenylyl cyclase activation, through several different mechanisms (reviewed in [56].) Mentioned above with regards to MMNSTs, *PRKAR1A* encodes an inhibitory subunit for protein kinase A. cAMP binds to PRKAR1A, which triggers conformational changes that dissociate PRKAR1A from the rest of the protein kinase A complex, releasing repression. Therefore, up-regulating protein kinase A activity might be a tumor promoting switch for both melanocytes and Schwann cells in dermal-like environments.

In the mouse uveal melanomas, almost all of the DE genes in *Nf1^flox^/+* tumors were unexpectedly related to muscle function. The down-regulated *COX6A* muscle gene was very significantly associated with worse uveal melanoma patient survival (p=1.7×10^−5^). *COX6A* encodes a subunit of the cytochrome c oxidase complex, the last enzyme in the mitochondrial electron transport chain. It’s down-regulation could therefore affect tumor metabolism. There are also some interesting muscle connections in the literature. For example, neurofibromin has been shown to be required for skeletal muscle development and function in mice [55] and individuals with neurofibromatosis type 1 experience hypotonia, decreased strength and reduced motor function [57]. Most intriguingly, people with myotonic dystrophy, which is caused by autosomal dominant repeat expansions disrupting the first exons of the muscle genes, *DMPK* (in type 1) or *CNBP* (in type 2), develop a significantly elevated number of thyroid, endometrium, ovary, melanoma, colon/rectum, and testis cancers [58]. One study found a 28-fold increased risk of uveal melanoma in people with myotonic dystrophy type 1 [59]. *CNBP* is located on chromosome 3, which is frequently lost in uveal melanoma. We checked the association between *DMPK* and *CNBP* expression and survival in the TCGA uveal melanoma dataset. Low *DMPK* expression was nearly associated with a worse outcome (p=0.054), and low *CNBP* expression also trended in that direction (p=0.16).

In summary, neurofibromin is a very large and complex tumor suppressor, the loss of which is known to contribute to the formation of melanoma in the skin and cause neurofibromas, a Schwann cell based tumor. Heterozygous 17q11.2 loss that includes the *NF1* locus is an uncommon, but recurrent phenomenon in intra-dermal and uveal melanoma that we think should be considered a potentially significant finding. In addition, our mouse model provides evidence that oncogenic GNAQ in post-natal *Plp1*-expressing cells causes nerve sheath like neoplasms, which should be further investigated.

## MATERIALS AND METHODS

### Mice

The research described in this article was conducted under the approval of the UBC Animal Care Committee (UBC animal care protocol number A19-0152, C.V.R). *Nf1^flox^* (*Nf1^tm1Par^*), *Plp1-cre ERT* (*Tg(Plp1-cre/ERT)3Pop*), *Rosa26-fs-GNAQ^Q209L^*(*Gt(ROSA)26Sor^tm1(GNAQ*)Cvr^*), and *Rosa26-fs*-*tdTomato* (*Gt(ROSA)26Sor^tm14(CAG-tdTomato)Hze^*) mice were genotyped as previously described [34,35,60,61,62]. Each allele was backcrossed to the C3HeB/FeJ genetic background for at least 4 generations before use. DNA from ear notches was isolated using the DNeasy Blood and Tissue kit (Qiagen) and amplified using PCR with HotStar Taq (Qiagen). Mice in the study were bred in two sequential cohorts. The first cohort established the development of tumors and the second cohort was used to increase numbers. There were roughly equal numbers of *Nf1^flox/^+* and *Nf1 +/+* mice expressing GNAQ^Q209L^ in each cohort.

### Tamoxifen

Tamoxifen (Sigma T5648) was dissolved in a corn oil/ethanol (10:1) mixture at a concentration of 10 mg/ml by gentle inversion at 37°C for 30 minutes, and then stored at 4°C for up to one week. At 5 weeks of age, sterile filtered tamoxifen (1 mg in 0.1 mL) was administered through intraperitoneal injection twice per day (every 12 hours) for 3 consecutive days.

### Animal monitoring

Animals were monitored every week to calculate a Clinical Health Score (CHS) in each of the following categories: weight, activity level, appearance, posture/gait, and tumor size (further details are available on request). Mice were euthanized when an externally visible tumor reached > 0.5 cm in diameter or there was another health concern affecting animal welfare (piloerection, hunching, excessive scratching, severely thickened ear skin without a single tumor > 0.5 cm.)

### Histology

Embedding and H&E staining was performed by WaxIt Histology services (Vancouver BC). Skin samples and tumors were fixed in 10% buffered formalin overnight at room temperature with gentle shaking, then were dehydrated, cleared, embedded in paraffin and sectioned at 5 μm, before staining using standard H&E technique. Eye samples were prepared in the same way, except fixation was in Davidson’s fixative for3 hours, followed by 10% formalin for 1 hour at 4°C. For visualization of Tomato fluorescence, eyes were fixed in 10% buffered formalin overnight at 4°C, taken through a sucrose gradient, embedded in O.C.T., and sectioned at 10 μm. Sections were washed in 1× PBS and counterstained with Dapi. Images were collected using an Axio Scan.Z1 slide scanner (Zeiss).

### Immunohistochemistry (IHC)

Our complete protocol for IHC on pigmented tissue will be submitted to *Bio-protocol* (https://bio-protocol.org/en). Briefly, 5 μm paraffin sections were first de-waxed and rehydrated into 1× PBS. Antigen retrieval was performed by incubating the slides in 0.6 L of 98°C citrate buffer pH 6.0 (Vector, H-3300) for 10 min, followed by removal from the heat source and cooling for 40 min at room temperature. Next, sections were washed and then placed in a solution of 10% H_2_O_2_ in 1× PBS, which was then heated to 60°C in an oven and held until all pigment was removed (2.5 hours). Sections were washed again, blocked in 5% normal serum in 1× PBS Triton 0.3% for 1 hr at room temperature and then incubated with the primary antibody (anti-S100b diluted 1:1000, Abcam, ab5264 or anti-RPE65 diluted 1:250, Thermo Fisher Scientific, MA1-16578; anti-CD34 diluted 1:200, abcam, ab8158) diluted in blocking solution for 1.5 hours at room temperature. Primary antibody was detected using an Elite ABC kit for HRP (Vector) using DAB with nickel (Vector, SK-4100) as directed. Some slides were then counterstained with haematoxylin. Stained slides were scanned for digital imaging (Panoramic 250 Flash III whole-slide scanner, 3DHISTECH). The sparse pigment tumors were incubated with myelin basic protein antibody (Abcam, ab218011) as above, but without the bleaching step, which was not compatible with this antibody.

### Measurement of spinal and CNS lesions

To quantify the surface area of the spine or brain that was affected, tissues were photographed under constant conditions alongside a ruler. In ImageJ, each cross-section image was transformed into a binary image. A freehand region of interest was drawn (around the lesion) and the threshold function was used to set threshold limits that covered the pigmented part of this area. The measure function was then used to compute the affected area in relative units. These were converted to mm^2^ using the ruler in each photo.

### Measurement of uveal melanomas

We computed the pigmented uveal tract thickness by analyzing eye H&E cross-sections using ImageJ. Images were taken from the center of the eye with the optic nerve present at the same magnification. A freehand region of interest was drawn (the eye) and the threshold function was used to set threshold limits that covered the pigmented tissue in the eye. The measure function was then used to compute this area in relative units. These were converted to mm^2^ using the scale bar in each image.

### RNA sequencing and DE analysis

All of the samples came from the 25 mice described in the text, except for 1 of the sparse pigment tumors, which came from a third round of tamoxifen injections to obtain another tumor of this type for RNAseq. Following euthanasia, tissue was collected from skin and eye tumors and placed in Trizol (LifeTechnologies) for homogenization. For skin, overlying epidermis and fat were removed before RNA extraction. For eyes, the globes were removed with curved forceps and placed on a chilled Petri dish on ice. Each eye was pulled open using fine forceps and pieces of thickened pigmented tissue were collected into the Trizol. RNA was isolated according to the manufacturer’s protocol. Further steps were performed by the BRC sequencing core at UBC. Quality control of the RNA samples was performed using the Agilent 2100 Bioanalyzer. Samples were then prepped according to the standard protocol for NEB next Ultra ii Stranded mRNA (New England Biolabs). All samples had an RNA Integrity Number (RIN) greater than or equal to 7.9. The samples were all poly(A) selected. RNA-sequencing was performed on the Illumina NextSeq 918 500 with Paired End 43bp x 43bp reads. The library consisted of ∼20-25 million reads in total per sample. Illumina’s bcl2fastq2 was used for demultiplexing the sequencing data and these reads were then aligned to the reference genome of Mus Musculus using STAR aligner. The FASTQ files are available at the Sequence Read Archive (SRA) database under project PRJNA1100739, (https://www.ncbi.nlm.nih.gov/sra/PRJNA1100739). The aligned read counts were used as input files for Differential Expression (DE) analysis using DESeq2 on R 4.0.3 following the “rnaseqGene” Bioconductor package.

### Statistical Analysis

Analyses for Gene ontology were performed using Enrichr. Statistical analysis of mouse phenotypes and survival were performed with Prism. TCGA-UVM Kaplan Meier survival curves were produced using Survival Genie [63].

## ACKNOWLEDGEMENTS

Funding for this research was provided by a grant from the Canadian Institutes of Health Research (CIHR), PJT-178178, to C.D. Van Raamsdonk.

## SUPPLEMENTARY FIGURE LEGENDS

**Supplementary Figure 1.**
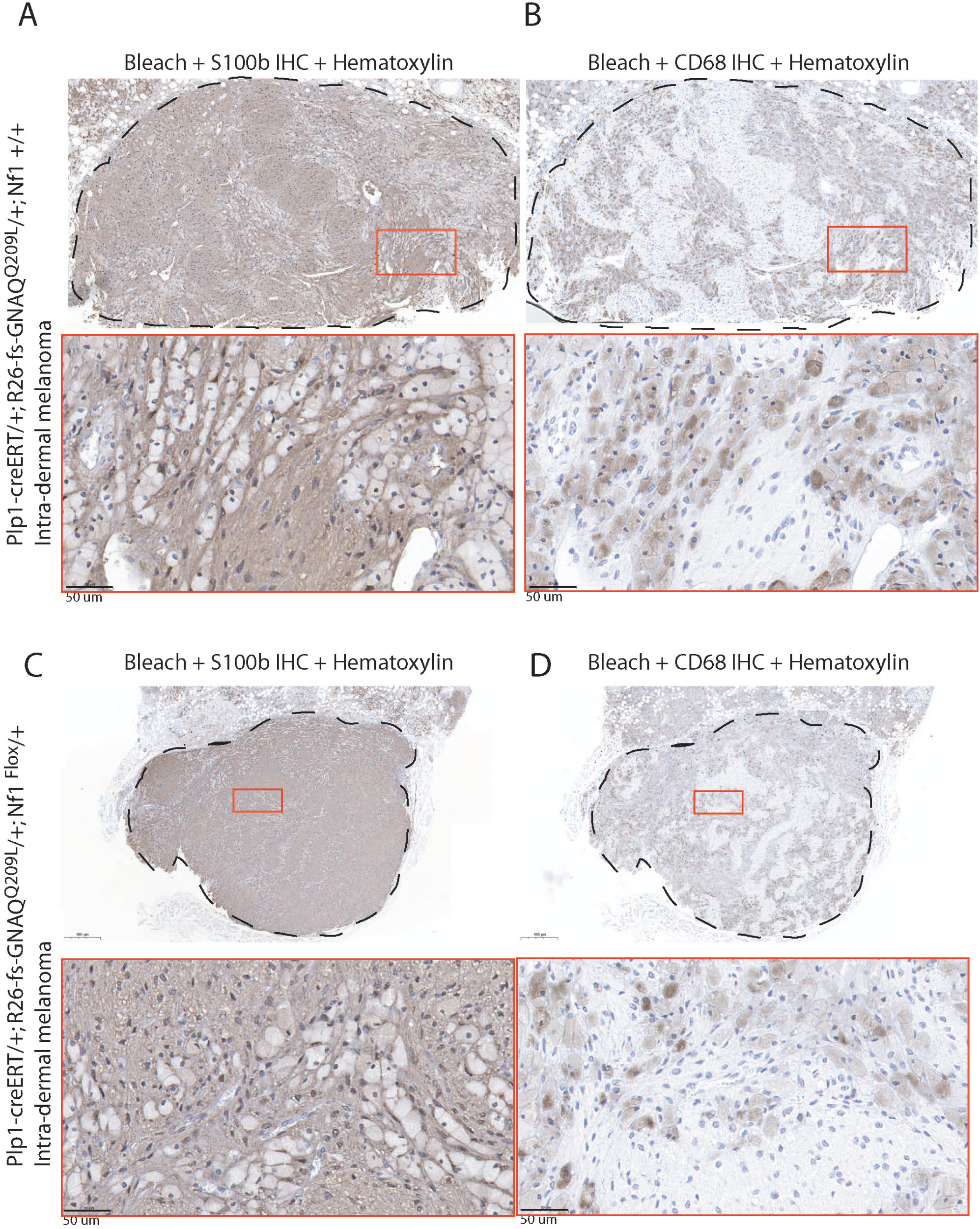
Histological analysis of intra-dermal melanomas. Representative images of S100b IHC (A,C) or CD68 (B,D) on intra-dermal melanomas from *Plp1-creERT*/+; *R26-fs-GNAQ^Q209L^*/+; +/+ (A,B) and *Plp1-creERT*/+; *R26-fs-GNAQ^Q209L^*/+; *Nf1^flox^*/+ (C,D) mice injected with tamoxifen at 5 weeks of age. Sections were bleached of melanin before antibody staining and counterstained with haematoxylin. In both tumors, there are a large number of CD68 positive cells, which have the cellular morphology of macrophages/melanophages (large, round, and prior to bleaching, pigmented). Note the complimentary patterns of S100b-positivity and CD68-positivity in the serial sections.

**Supplementary Figure 2.**
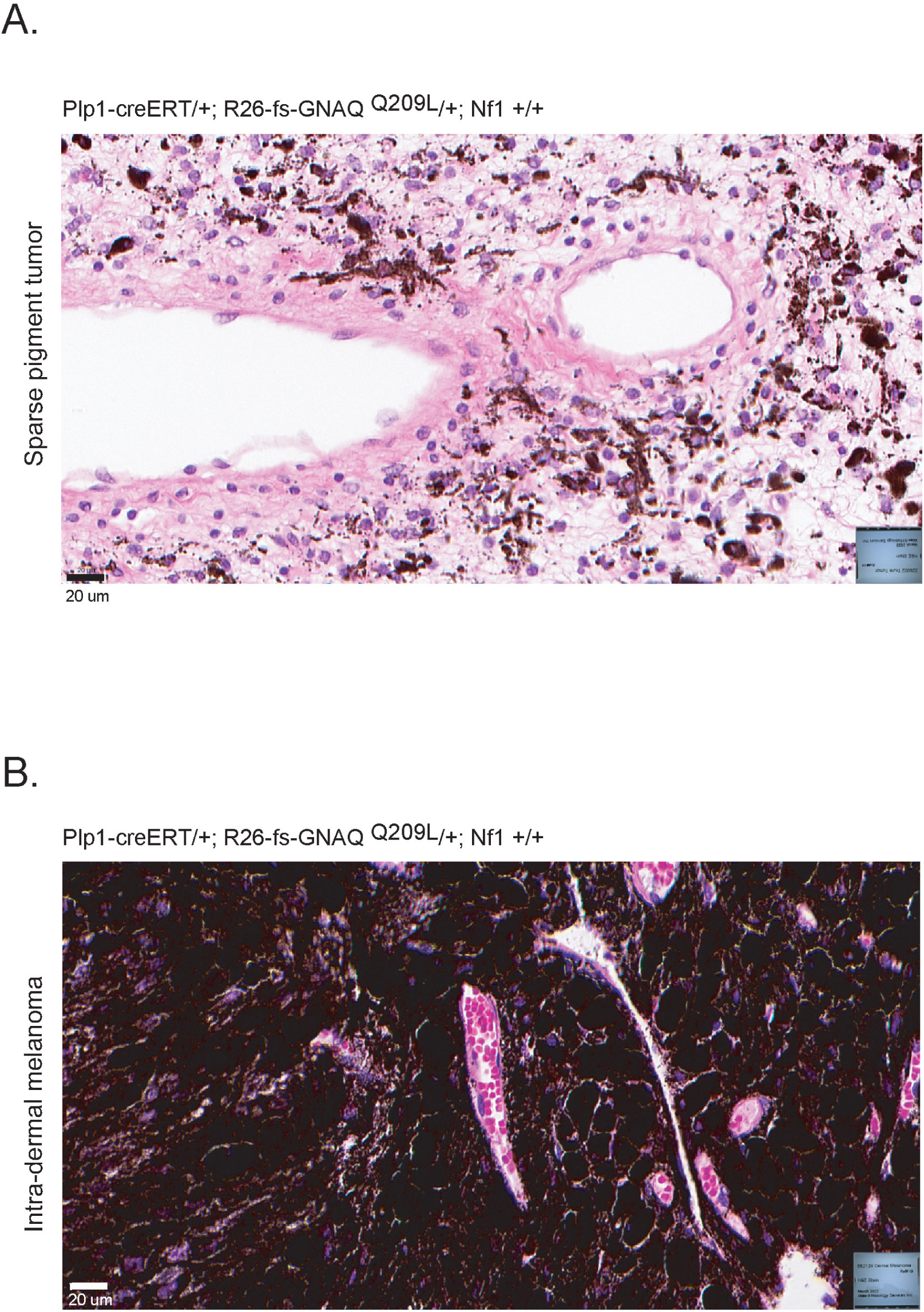
Comparison of blood vessels in a sparse pigment tumor and an intra-dermal melanoma from *Plp1-creERT*/+; *R26-fs-GNAQ^Q209L^*/+; +/+ mice. **(A,B)** H&E stained sections of a sparse pigment tumor (A) and an intra-dermal melanoma (B). The vessel walls of the sparse pigment tumor are thicker and stained brighter pink with eosin.

**Supplementary Figure 3.**
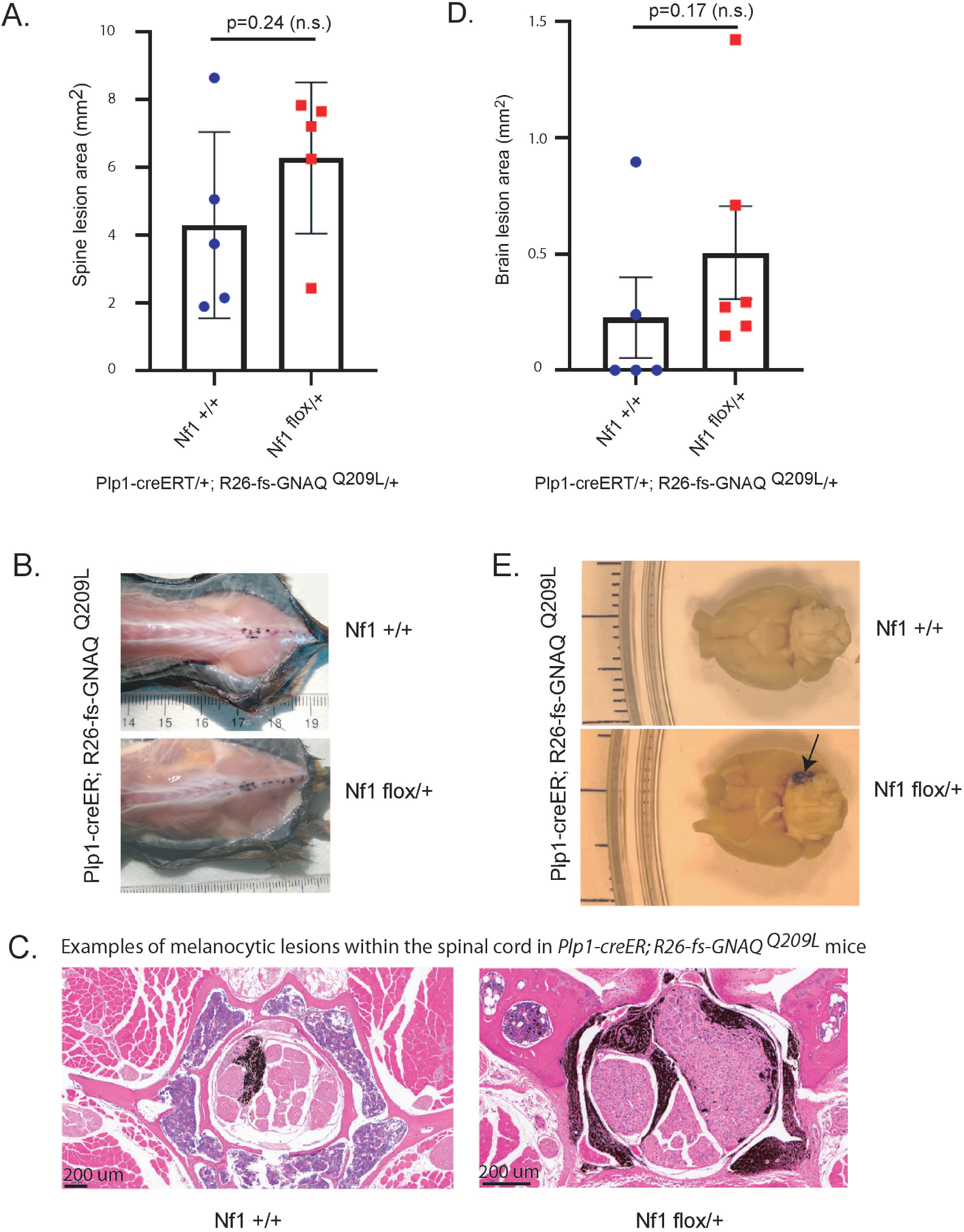
There was no significant difference found in CNS phenotypes between *Nf1* genotypes. **(A)** Graph showing average area of pigmented lesions associated with the spine in *Plp1-creERT*/+; *R26-fs-GNAQ^Q209L^*/+; *Nf1^flox^*/+ and *Plp1-creERT*/+; *R26-fs-GNAQ^Q209L^*/+; +/+ mice injected with tamoxifen at 5 weeks of age and euthanized upon tumor or other humane endpoint. **(B)** Representative *Plp1-creERT*/+; *R26-fs-GNAQ^Q209L^*/+; *Nf1^flox^*/+ and *Plp1-creERT*/+; *R26-fs-GNAQ^Q209L^*/+; +/+ mice exhibiting darkly pigmented lesions associated with the spine. (**C)** H&E stained sections of spines showing melanocytic growth within neural tissues. **(D)** Graph showing average area of pigmented lesions associated with the ventral brain surface in *Plp1-creERT*/+; *R26-fs-GNAQ^Q209L^*/+; *Nf1^flox^*/+ and *Plp1-creERT*/+; *R26-fs-GNAQ^Q209L^*/+; +/+ mice. **(E)** Representative *Plp1-creERT*/+; *R26-fs-GNAQ^Q209L^*/+; *Nf1^flox^*/+ and *Plp1-creERT*/+; *R26-fs-GNAQ^Q209L^*/+; +/+ brains, ventral side up. Arrow indicates a pigmented lesion. Although there was no significant difference, the trends suggest that with more animals, *Nf1^flox^/+* might cause greater growth of CNS associated lesions. Error bars in graphs represent the standard error of the mean (SEM).

**Supplementary Figure 4.**
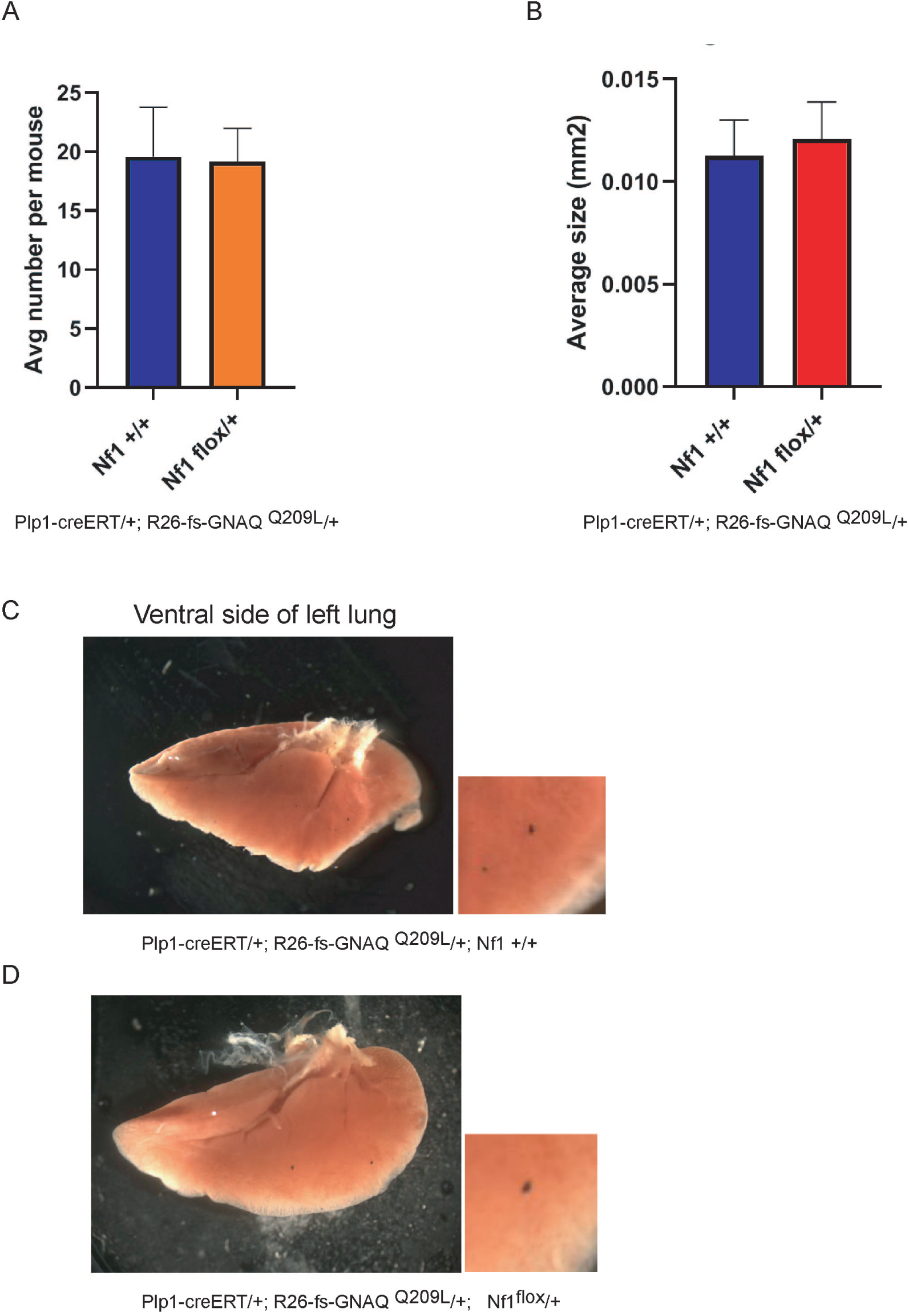
There was no significant difference in lung phenotypes between *Nf1* genotypes. **(A,B)** Graphs showing average number of pigmented lesions and the average size of pigmented lesions in the lungs of *Plp1-creERT*/+; *R26-fs-GNAQ^Q209L^*/+; +/+ or *Plp1-creERT*/+; *R26-fs-GNAQ^Q209L^*/+; *Nf1^flox^*/+ mice injected with tamoxifen at 5 weeks of age and euthanized upon tumor or other humane endpoint. **(C,D)** Representative left lungs from *Plp1-creERT*/+; *R26-fs-GNAQ^Q209L^*/+; +/+ (C) and *Plp1-creERT*/+; *R26-fs-GNAQ^Q209L^*/+; *Nf1^flox^*/+ (D) mice. Error bars in graphs represent the standard error of the mean (SEM).

**Supplementary Figure 5.**
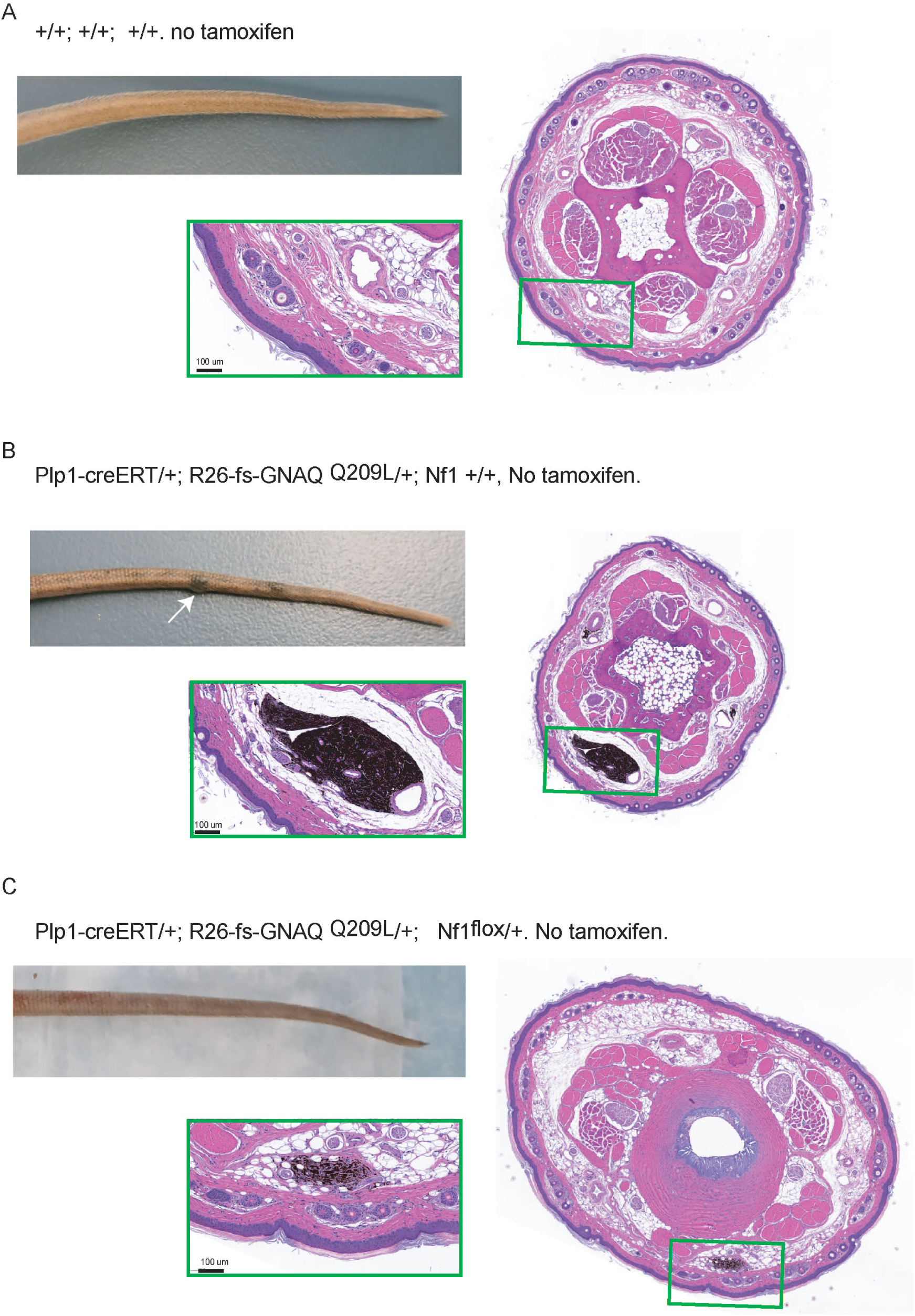
Tail phenotypes in control mice not injected with tamoxifen. **(A,B,C).** Whole tails and H&E stained sections of +/+; +/+; +/+ (A), *Plp1-creERT*/+; *R26-fs-GNAQ^Q209L^*/+; +/+ (B), or *Plp1-creERT*/+; *R26-fs-GNAQ^Q209L^*/+; *Nf1^flox^*/+ (C) mice that were housed in tamoxifen-free cages. Two cases of tail lesions that were found are shown in B and C, indicating that there is some small amount of leaky CreERT activity. Animals were healthy and aged to 72 weeks.

**Supplementary Figure 6.**
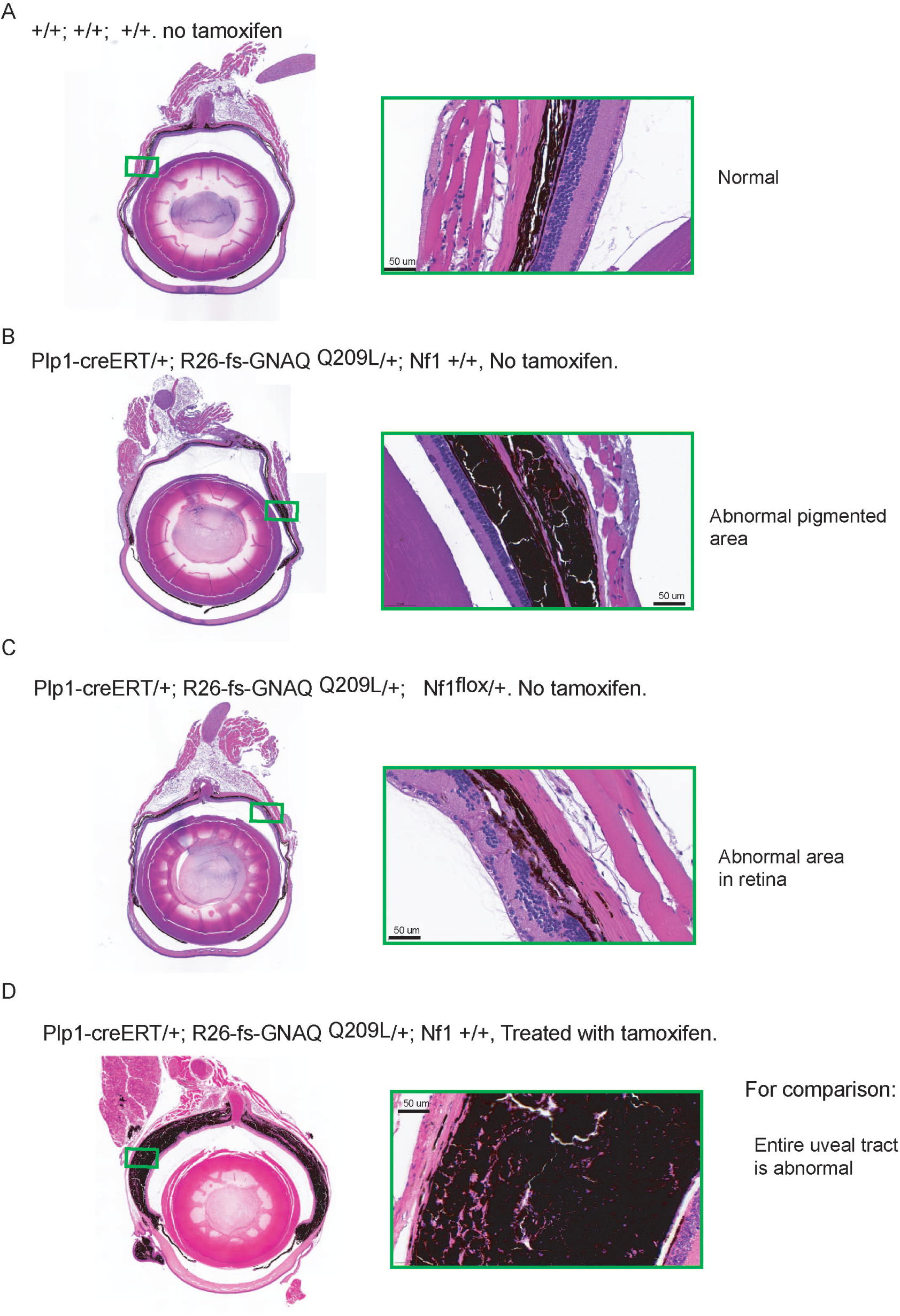
Eye phenotypes in control mice not injected with tamoxifen. **(A,B,C).** Eye sections stained with H&E from +/+; +/+; +/+ (A), *Plp1-creERT*/+; *R26-fs-GNAQ^Q209L^*/+; +/+ (B), or *Plp1-creERT*/+; *R26-fs-GNAQ^Q209L^*/+; *Nf1^flox^*/+ (C) mice that were housed in tamoxifen-free cages until 72 weeks old. One eye was found with a small area of expanded pigmentation (boxed area in B), which suggests that there is some small amount of leaky CreERT activity in ocular melanocytes. Another eye exhibited a disorganized neural retina, of unknown significance to the study (boxed area in C). **(D)** For reference, a tamoxifen treated *Plp1-creERT*/+; *R26-fs-GNAQ^Q209L^*/+; +/+ mouse eye shows a much more enhanced growth of the uveal tract.

**Supplementary Figure 7.**
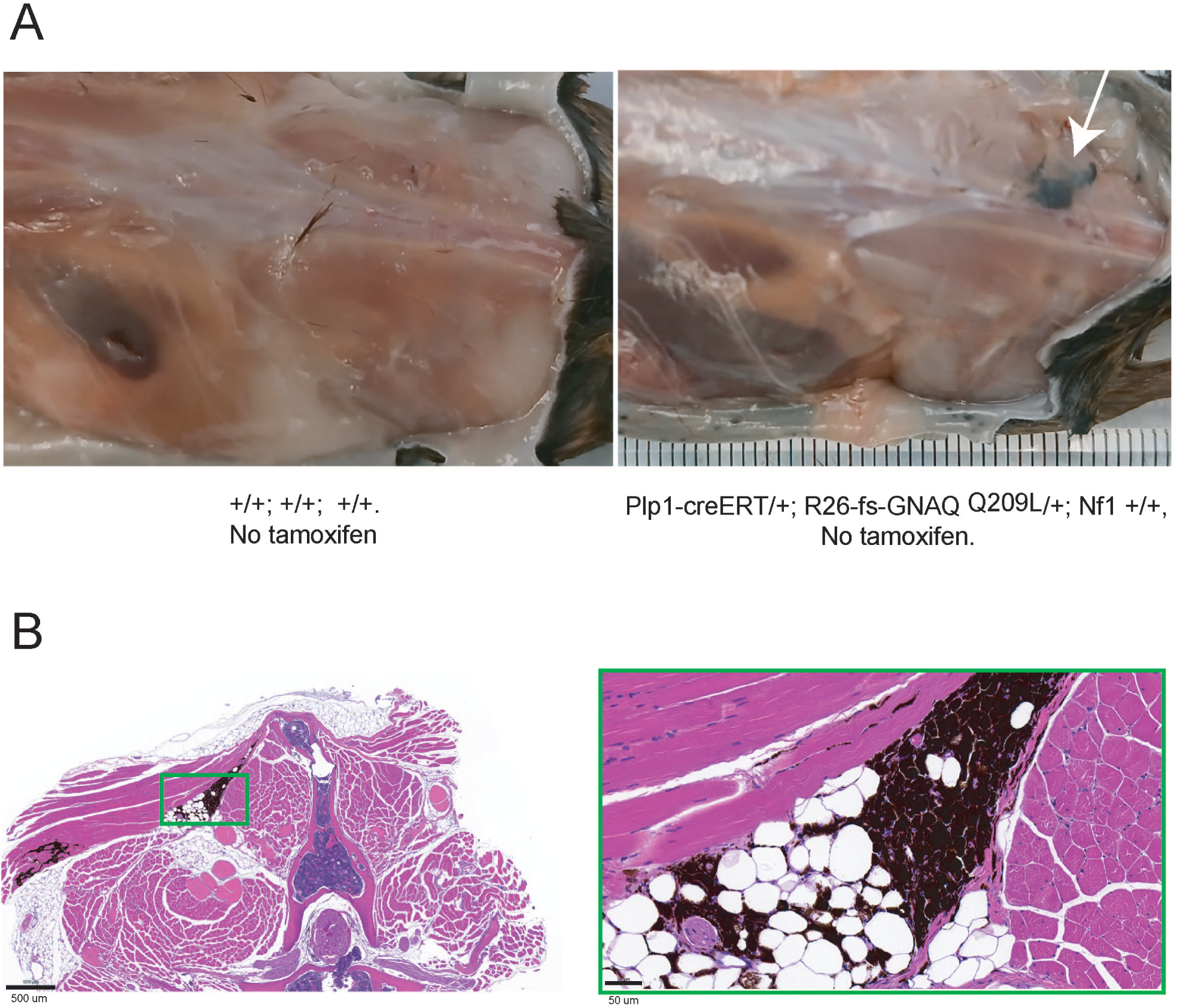
Spine phenotypes in control mice not injected with tamoxifen. **(A)**. Macroscopic image of the spine of a+/+; +/+; +/+ (left) and *Plp1-creERT*/+; *R26-fs-GNAQ^Q209L^*/+; +/+ (right) mouse housed in tamoxifen-free cages until 72 weeks old. The *Plp1-creERT*/+; *R26-fs-GNAQ^Q209L^*/+; +/+ mouse exhibited a pigmented lesion, indicated with the arrow. **(B)** H&E stained section through the lesion shown in A. Invasion of the muscle layer is apparent. This lesion suggests that there is some small amount of leaky CreERT activity in melanocytes associated with the CNS and normally found in the meninges.

## SUPPLEMENTARY TABLE LEGENDS

**Supplementary Table 1a:** Survival data in tamoxifen injected mice expressing GNAQ^Q209L^

**Supplementary Table 1b:** Results of differential gene expression analysis comparing *Nf1 ^flox^*/+ versus *Nf1* wildtype intra-dermal melanomas.

**Supplementary Table 1c:** Significant p-values and q-values for gene ontology analysis of DE genes in *Nf1 ^flox^*/+ intra-dermal melanomas.

**Supplementary Table 1d:** Results of differential gene expression analysis comparing *Nf1 ^flox^*/+ versus *Nf1* wildtype uveal melanomas.

**Supplementary Table 1e:** Significant p-values and q-values for gene ontology analysis of DE genes in *Nf1 ^flox^*/+ uveal melanomas.

**Supplementary Table 1f:** Results of differential gene expression analysis comparing sparse pigment tumors versus intra-dermal melanomas.

**Supplementary Table 1g:** Significant p-values and q-values for gene ontology analysis for cell type in sparse pigment tumors.

## Notes

https://www.ncbi.nlm.nih.gov/sra/PRJNA1100739

### Competing Interest Statement

The authors have declared no competing interest.

